# Heterogeneity and Functional Analysis of Cardiac Fibroblasts in Heart Development

**DOI:** 10.1101/2023.07.30.551164

**Authors:** Yiting Deng, Yuanhang He, Juan Xu, Haoting He, Guang Li

**Affiliations:** Tsinghua University, School of Medicine; Department of Developmental Biology, University of Pittsburgh School of Medicine, Pittsburgh, PA 15201, USA

**Keywords:** Heart development, Single cell, Fibroblast, Heterogeneity, Ablation, Extracellular matrix, Ligand-receptor

## Abstract

**Background:** As one of the major cell types in the heart, fibroblasts play critical roles in multiple biological processes. Cardiac fibroblasts are known to develop from multiple sources, but their transcriptional profiles have not been systematically compared. Furthermore, while the function of a few genes in cardiac fibroblasts has been studied, the overall function of fibroblasts as a cell type remains uninvestigated.

**Methods:** Single-cell mRNA sequencing (scRNA-seq) and bioinformatics approaches were used to analyze the genome-wide genes expression and extracellular matrix genes expression in fibroblasts, as well as the ligand-receptor interactions between fibroblasts and cardiomyocytes. Single molecular in situ hybridization was employed to analyze the expression pattern of fibroblast subpopulation-specific genes. The Diphtheria toxin fragment A (DTA) system was utilized to ablate fibroblasts at each developmental phase.

**Results:** Using RNA staining of *Col1a1* at different stages, we grouped cardiac fibroblasts into four developmental phases. Through the analysis of scRNA-seq profiles of fibroblasts at 18 stages from two mouse strains, we identified significant heterogeneity, preserving lineage gene expression in their precursor cells. Within the main fibroblast population, we found differential expressions of Wt1, Tbx18, and Aldh1a2 genes in various cell clusters. Lineage tracing studies showed Wt1- and Tbx18-positive fibroblasts originated from respective epicardial cells. Furthermore, using a conditional DTA system-based elimination, we identified the crucial role of fibroblasts in early embryonic and heart growth, but not in neonatal heart growth. Additionally, we identified the zone- and stage-associated expression of extracellular matrix genes and fibroblast-cardiomyocyte ligand-receptor interactions. This comprehensive understanding sheds light on fibroblast function in heart development.

**Conclusion:** We observed cardiac fibroblast heterogeneity at embryonic and neonatal stages, with preserved lineage gene expression. Ablation studies revealed their distinct roles during development, likely influenced by varying extracellular matrix genes and ligand-receptor interactions at different stages.

## Introduction

Fibroblasts are one of the most abundant cardiac cell types and play important roles in normal heart function and pathological heart remodeling at the adult stage^1, 2^. The fibroblast composition in the heart has been analyzed with multiple approaches such as flow cytometry, with a focus on the left ventricle^3, 4^. However, the anatomical location of fibroblasts in the entire heart, including the atrial chambers, along the developmental progression is still unclear. The heart in mice starts to develop into a four-chambered structure at E9.5. The atrial and ventricular chambers are connected by the atrioventricular canal (AVC), a transient structure that develops into the septum and valve cells^5, 6^. Although the main fibroblast population does not develop until E13.5^7^, fibroblast-like cells with the expression of *Col1a1* and other fibroblast genes start to develop in the AVC at E9.5^6^.

Cardiac fibroblasts have been reported to develop from multiple sources^8^. The epicardial cell is the main source and contributes to most resident fibroblasts. The endocardial endothelial cell contributes to the fibroblasts in the ventricular septum and part of the fibroblasts in the ventricular chambers. Neural crest cells also contribute to cardiac fibroblast development. Lineage tracing results showed that they mainly differentiate into fibroblasts in large vessels and the right atrium^8–10^. The fibroblasts in valves (valve interstitial cells) are developed from endothelial cells and epicardial cells in atrioventricular valves, and from endothelial cells and neural crest cells in semilunar valves^11^.

The epicardial cell, the main source of fibroblasts, is located on the outer surface of the heart and develops from the proepicardium^12^. Epicardial cells have been found to contribute to several cell types in the heart, such as fibroblasts and smooth muscle cells^13^. They express many marker genes, such as Wt1, Tbx18, Aldh1a2, Tcf21, and others^14^. The heterogeneity of epicardial cells has been observed in different contexts, such as heart regeneration. In zebrafish, transient epicardial cell subpopulations expressing Hapln1 or Ptx3a have been observed after injury and play important roles in heart regeneration^15, 16^. However, it is still elusive whether there is heterogeneity in epicardial cells during normal heart development. Although no epicardial cell subpopulations were identified before E13.5 through RNA staining of multiple epicardial genes^17^, it is unknown whether epicardial cells develop subpopulations at later stages and contribute to different lineage descendants.

The main function of fibroblasts in tissue development is to synthesize extracellular matrix (ECM) proteins and secrete growth factors that influence the development of other lineages^2, 18^. Using a cell coculture system, embryonic cardiac fibroblasts-derived ECMs, such as collagen and fibronectin, were found to promote cardiomyocyte proliferation through β1-integrin signaling^19^. However, their in vivo function remains unclear. To study the function of a cell type rather than a gene, toxin genes such as NTR and DTA have been engineered into conditional ablation systems to eliminate specific groups of cells^20, 21^. The DTA system has been used in mice to ablate cardiac progenitor cells and cardiomyocytes at early developmental stages^22^. Recently, a group of proliferating fibroblasts with the expression of Postn was identified in neonatal hearts. Further elimination of this cell population using the DTA ablation system showed its importance in promoting cardiomyocyte maturation^23^.

Over the years, single-cell mRNA sequencing (scRNA-seq) has proved its power in dissecting tissue heterogeneity and identifying associated gene expression profiles. Cardiac fibroblasts, along with fibroblasts in several other tissues in adult mice, were profiled using scRNA-seq and revealed tissue-specific gene expression signatures^24^. These signatures were then used to guide the in vitro generation of cardiac fibroblasts from human stem cells^25^. Recently, our lab has generated a scRNA-seq atlas covering all the main cardiac cell types, including fibroblasts at embryonic and neonatal stages^26^. This atlas serves as a valuable resource to further explore the heterogeneity of each cell type, including fibroblasts, and their interactions with other cell types.

In this study, we analyzed the cardiac fibroblast scRNA-seq data at embryonic and neonatal stages and identified heterogeneities at multiple levels. We identified four distinct populations and several subpopulations within the main population. We further used lineage tracing and RNA staining to show that the subpopulations were derived from different epicardial cells. Additionally, we identified four phases of cardiac fibroblasts according to their developmental progression. Using a DTA-based cell ablation system, we eliminated the fibroblasts at each phase and observed different outcomes. Finally, through the analysis of extracellular matrix genes expression in fibroblasts and ligand-receptor interactions between fibroblasts and cardiomyocytes, we identified chamber- and stage-associated features that probably contributed to the differential ablation outcomes. In summary, this study provides a comprehensive understanding of the heterogeneity and function of cardiac fibroblasts at embryonic and neonatal stages.

## Results

### RNA staining revealed the anatomical location of cardiac fibroblasts at different stages

To understand the development of cardiac fibroblasts (FBs) in terms of their spatial distributions along the developmental progression, we stained the fibroblast marker gene *Col1a1* on heart sections at different stages, including E11.5, E14.5, E16.5, E17.5, P2, and P3. We observed a strong expression of *Col1a1* in epicardial cells at all analyzed stages (Figure 1A). Within the heart, we found *Col1a1*-positive cells enriched at the boundary of the atrial and ventricular regions in E11.5 hearts (Figure 1Ai), indicating their presence in the atrioventricular canal (AVC). By E14.5, the AVC cells had developed into valve structures, and the valve interstitial cells highly expressed *Col1a1* (Figure 1Aii). In the ventricles, a small proportion of *Col1a1*-positive cells were observed adjacent to the epicardium. In contrast, no obvious *Col1a1* signal was found in the atrium at this stage (Figure 1Aii). At E16.5, both ventricles were filled with *Col1a1*-positive cells, but the middle of the ventricular septum lacked signal (Figure 1Aiii). FBs were also found in the atrium, but the signal at the tips of both atrial chambers was sparser (Figure 1Aiii). The distribution of FBs at E17.5 was highly similar to that observed at E16.5 (Figure 1Aiv). Furthermore, we analyzed the *Col1a1* signal in P2 and P3 hearts and found dense *Col1a1* expression at the valves and in all four chambers (Figure 1Av, vi). In summary, this analysis revealed the spatial location of cardiac FBs along the developmental stages. Specifically, we found that the FBs developed first in the AVC and valve, then at the ventricular wall, and last at the ventricular septum (Figure 1B).

**Fig 1.**
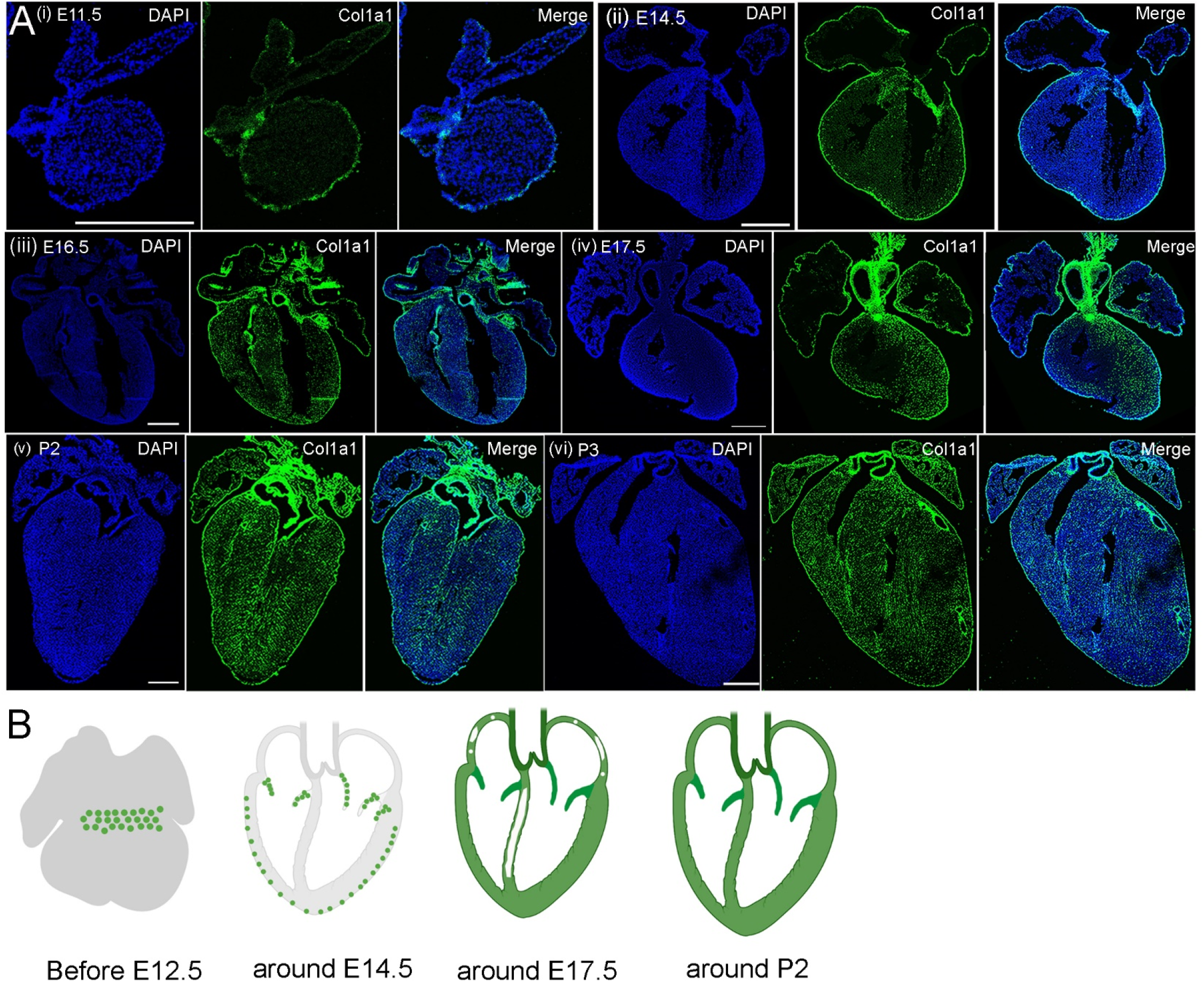
Anatomical location of cardiac fibroblasts at different stages. (Ai-vi) RNA staining of *Col1a1* revealed the spatial pattern of cardiac fibroblasts at different developmental stages. (B) The development of cardiac fibroblasts can be grouped into four phases. Scale bar=500µm.

### Identification of heterogeneity in cardiac fibroblasts through the analysis of scRNA-seq data

Through the analysis of single-cell RNA-sequencing (scRNA-seq) data at 18 stages of developing hearts from CD1 mice, we investigated the expression of *Col1a1* and observed high expression in epicardial cells and fibroblasts (Figure 2A). This is consistent with the findings from RNA staining (Figure 1A). Subsequently, we reanalyzed the fibroblasts and identified four distinct populations comprising 13 clusters (Figure 2D). Gene expression analysis revealed that all cells in these populations exhibited high expression of *Col1a1* (Figure 2A). Moreover, through differential gene expression analysis between the populations (Supplementary table S2), we discovered that one group of cells highly expressed *Hapln1* (Figure 2B, D, E), a marker gene associated with valves. This indicates that these cells are valve interstitial cells. We also found another group of cells expressing *Cdh5* and *Tie1* (Figure 2B, D, E), genes associated with the endothelial cell (EDC) lineage. This suggests that these cells may represent fibroblasts derived from EDCs. Additionally, we identified a third group of fibroblasts expressing *Sox10* and *Phox2b* (Figure 2B, D, E), genes associated with the neural crest cell lineage. This suggests that these cells may originate from neural crest cells. Finally, the rest of the cells formed a main population that likely represents epicardial cell-derived fibroblasts in the chambers, as they expressed *Tcf21* and other marker genes (Figure 2D, E). Interestingly, further analysis of the cell cycle phases in these fibroblasts revealed that each population consisted of cells in all three phases (Figure 2B), indicating active proliferation during the early stages.

**Fig 2.**
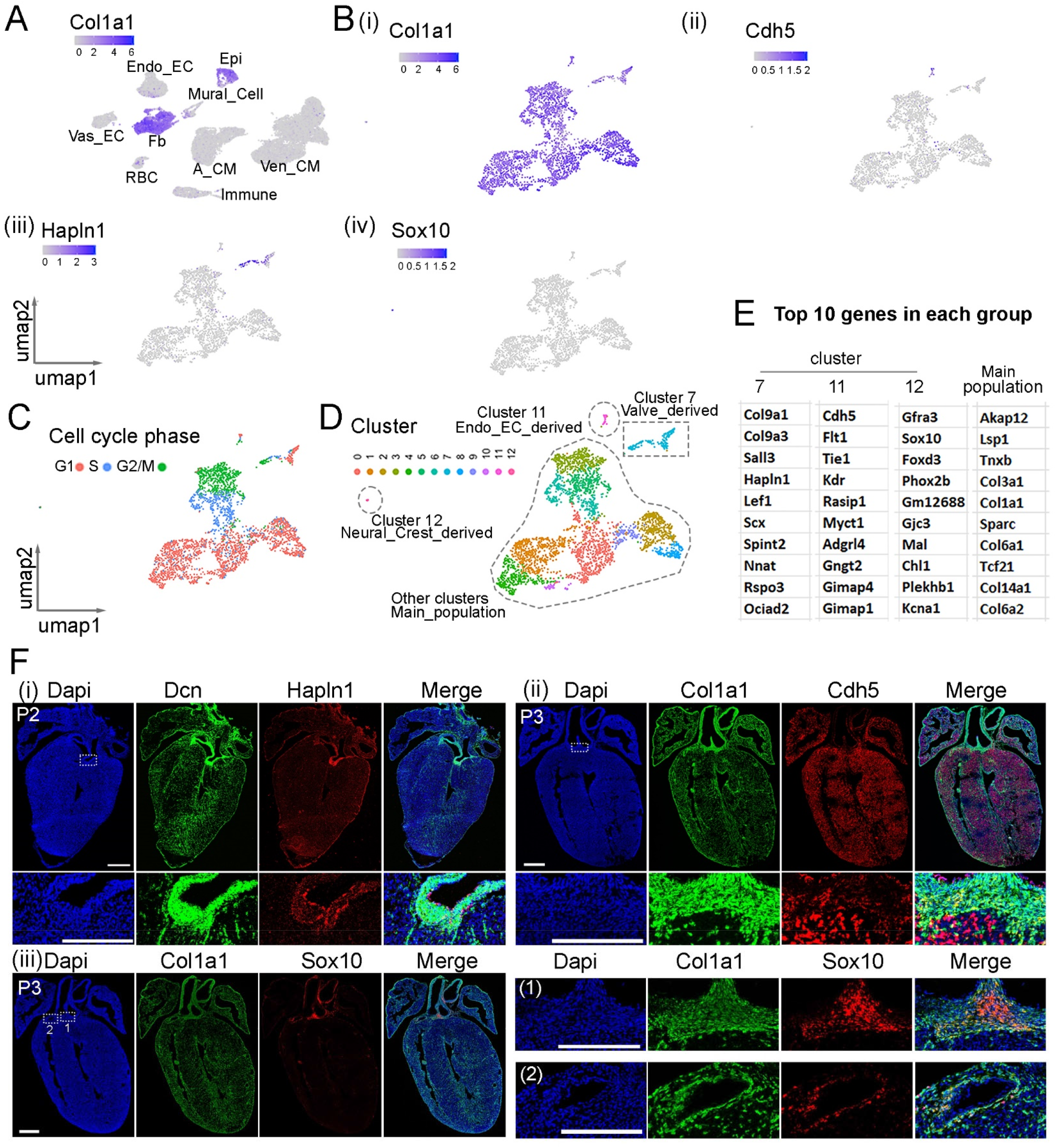
ScRNA-seq identified distinct fibroblast populations. (A) UMAP plot of *Col1a1* expression in cardiac cells. (Bi-iv) UMAP plots of *Col1a1* and representative cluster-specific genes expression in cardiac fibroblasts. (C) UMAP plot of fibroblasts labeled by cell cycle phases. (D) Diagram of the four types of cardiac fibroblasts. (E) The top 10 genes expressed in each group of cardiac fibroblasts. (Fi-iii) Identification of the anatomical location of each fibroblast population through RNA staining. Scale bar=500µm and 250µm in the whole heart images and the images with enlarged areas, respectively.

We also analyzed the heart scRNA-seq data from C57BL/6 mice and identified the fibroblasts with *Col1a1* expression (Supplementary table S3). Re-clustering analysis of the *Col1a1*-positive cells also revealed four populations comprising 11 clusters (Figure S1A, B, D). Differential gene expression analysis between populations revealed a Hapln1-positive population and a Sox10-positive population. However, we did not observe a Cdh5-positive population, which was probably missed due to a small proportion in the cardiac fibroblast population. In contrast, we identified a population of cells expressing Prdm6, a marker gene for semilunar valve cells (Figure S1A). Further phase analysis of the cells revealed that each population has cells from all cell cycle phases (Figure S1C). Next, we performed an integrative analysis of valve cells from CD1 and C57BL/6 mouse scRNA-seq datasets. The valve cells formed 7 clusters with the identities of different cell cycle phases, stages, and zones (Figure S2A-D). We also observed that Prdm6 and Hapln1 were expressed at different cell clusters (Figure S2E-F). Interestingly, we found that the Hapln1-positive cells were from both the left ventricle (LV) and right ventricle (RV), and the cells from both chambers did not form distinct clusters. This suggests that the fibroblasts from the mitral valve and tricuspid valve are transcriptionally similar. In contrast, we found that the Prdm6-positive cells were mostly from the RV, suggesting that the semilunar valve fibroblasts we profiled mostly were pulmonary valve cells. Moreover, through differential gene expression analysis between cell clusters, we found that the cluster 1 cells were mainly from RV and highly expressed Meox1 (Figure S2G), a gene that was found to be expressed in VICs on the inner side of aortic valve leaflets^27^. This suggests that the cluster of these cells may be located on the inner side of tricuspid valve leaflets.

Next, to understand the anatomical location of these fibroblast populations, we performed RNA in situ hybridizations on postnatal heart sections. Firstly, we stained *Hapln1* together with *Dcn*, another Pan-fibroblast marker gene. We found that *Hapln1* was highly expressed in valves, and most of its signal overlapped with *Dcn* (Fig 2Fi), suggesting its expression in valve interstitial cells (fibroblast-like cells in valves). We also stained *Cdh5* with *Col1a1* and observed a small cluster of cells with the expression of both genes, mostly located at the boundary of large vessels and ventricular chambers (Fig 2Fii). Finally, we co-stained *Sox10* and *Col1a1* expression (Fig 2Fiii). As previously reported^10^, we identified double positive cells in the outflow tract-derived large vessels (Fig 2Fiii_1). However, interestingly, we also observed their presence in coronary vessels (Fig 2Fiii_2), suggesting that these cells may have a broader function than previously thought. In summary, through the analysis of scRNA-seq data and in situ hybridizations, we identified different populations of cardiac FBs, which differentially expressed lineage genes likely from their precursor cells.

### Subpopulations in the main fibroblast population exhibited differential expression of epicardium lineage genes

Subsequently, we reanalyzed the fibroblasts in the main population. Stage analysis of these fibroblasts revealed a spatial pattern where cells in early stages occupied the middle and gradually transitioned towards fibroblasts in later stages in two directions (Fig 3Ai). Further analysis of the cell cycle phases indicated that the fibroblasts on the left were in the G1 phase, while those on the right were in the S and G2/M phases (Fig 3Aiii). Employing unsupervised clustering analysis, we grouped the fibroblasts into eight clusters, consisting of six clusters of G1 cells (clusters 1, 2, 3, 5, 6, 7) and two clusters of S/G2/M cells (clusters 0, 4) (Fig 3C). Additionally, differential gene expression analysis revealed that the cells in the G1 phase exhibited differential expression of three epicardial cell lineage genes (Fig 3B, Supplementary table S4). Specifically, we observed that cluster 7 expressed Wt1 and Tbx18, clusters 1 and 6 expressed Aldh1a2, cluster 5 expressed Wt1, cluster 2 expressed Tbx18 and Aldh1a2, and cluster 3 expressed Tbx18 only (Fig 3B, C). After considering the source chamber and stage for the cells in each cluster, we found that the Wt1 and Tbx18 double positive FBs (C7) were mainly from the stages before E14.5 without obvious chamber preferences (Fig 3B, C, S3Ai). Interestingly, in the G2/M phased cells, we found that cluster 4 cells with the expression of Wt1 and Tbx18 were also mainly from the stages before E14.5 (Fig 3B, C, S3Bii), suggesting that the FBs at early stages expressed Wt1 and Tbx18 regardless of their cell cycle status. For the cells with Aldh1a2 expression only (C1_6), we found that they were mainly from ventricular at stages after E13.5 (Fig 3B, C, S3Aiv). In contrast, the Wt1 positive cells (C5) were found to be mainly from the atrium at stages after E13.5. Additionally, we did not observe a Wt1 positive cell population in the G2/M phase at the same period (Fig 3B, C, S3Aii), suggesting that this cell population was less proliferative. In the atrium, we identified another two cell clusters at later stages. The cluster 2 cells with the expression of Tbx18 and Aldh1a2 were mainly from RA, and the cluster 3 with Tbx18 expression was mainly from LA (Fig 3B, C, S3Aiii, v). The cluster 0 with G2/M phased cells is a mixture of Aldh1a2 positive cells, Tbx18 positive cells, and their double positive cells from stages after E13.5 (Fig 3A, B, S3Bi). Finally, we conducted a similar analysis on the main FB population from C57BL/6 mice and observed the same five groups of FBs (Fig S4A-E, Supplementary table S5).

**Fig 3.**
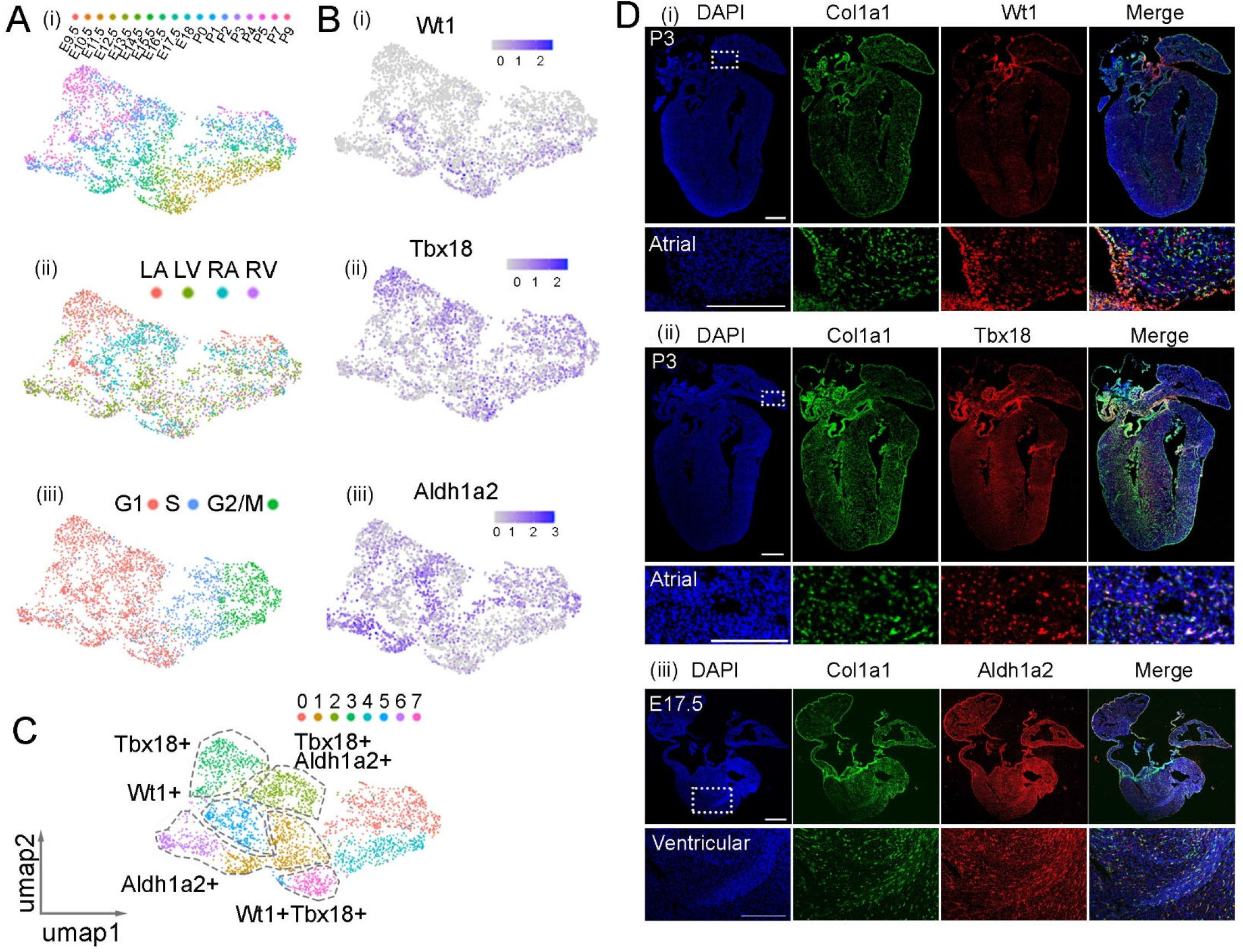
Identification of subpopulations in the epicardium-derived fibroblast (Epi_FB) population. (Ai-iii) UMAP plots of Epi_FBs labeled by stage, zone, and cell cycle phases. (B) UMAP plots showing the expression of *Wt1*, *Tbx18*, and *Aldh1a2*. (C) Identification of five subpopulations in Epi_FB that differentially expressed *Wt1*, *Tbx18*, and *Aldh1a2*. (Di-iii) RNA staining analysis of *Col1a1* together with *Wt1*, *Tbx18*, or *Aldh1a2*. Scale bar = 500µm for the whole heart section, and 250µm for the enlarged sections.

To validate these subpopulations, we performed RNA staining of *Col1a1* together with *Wt1*, *Tbx18*, or *Aldh1a2* on heart sections at later developmental stages. Specifically, we analyzed *Wt1* and *Tbx18* at E17.5 and P3 using RNAScope and PLISH. For *Aldh1a2* staining, we only performed it at E17.5 with PLISH. Consistent with scRNA-seq data, we observed Wt1 and *Tbx18* positive fibroblasts mainly in the atrial region, while *Aldh1a2* positive fibroblasts were predominantly located in the ventricular region (Fig 3Di-iii). Interestingly, we found that the *Tbx18* positive fibroblasts were mainly situated in the middle of the atrial region, while *Wt1* positive fibroblasts were predominantly found at the edges (Fig 3Di, ii, S5A-D). This suggests that the *Wt1* positive cells may be newborn cells derived from epicardial epithelial-mesenchymal-transition (EMT).

### Lineage analysis of fibroblast subpopulations

To understand if the *Wt1* and *Tbx18* positive fibroblasts were developed from epicardial cells expressing the same genes, we performed lineage tracing by breeding two mouse strains, Wt1-CreER and Tbx18-CreER, with the reporter mouse line Rosa26-mTmG. Since fibroblasts in the atrial region have not yet developed at E14.5, we treated pregnant dams with tamoxifen at both E13.5 and E14.5 to label the epicardial cells (Fig 4A, D). For the Wt1-CreER;mTmG mice, we harvested the labeled hearts at P0 and performed RNAScope to analyze the expression of *Wt1* or *Tbx18* together with *Col1a1* and *eGFP*. Interestingly, we found that the *Wt1* and *Col1a1* positive cells at the edge of the atrial region were also *eGFP* positive, suggesting that these cells were derived from *Wt1* positive precursors. In contrast, the *Tbx18* and *Col1a1* positive cells in the middle of the atrial region were *eGFP* negative, indicating that these fibroblasts were not developed from *Wt1* positive cells (Fig 4B, C). Next, we performed the same analysis on Tbx18-CreER;mTmG labeled hearts at E19.5. We found that the fibroblasts in the middle of the atrial region were *Tbx18* and *eGFP* positive but not Wt1 positive (Fig 4E, F), suggesting that the *Tbx18* positive fibroblasts were derived from *Tbx18* positive precursors. In summary, the lineage tracing results indicated that *Wt1* and *Tbx18* positive fibroblasts were derived from *Wt1* and *Tbx18* positive epicardial cells, respectively.

**Fig 4.**
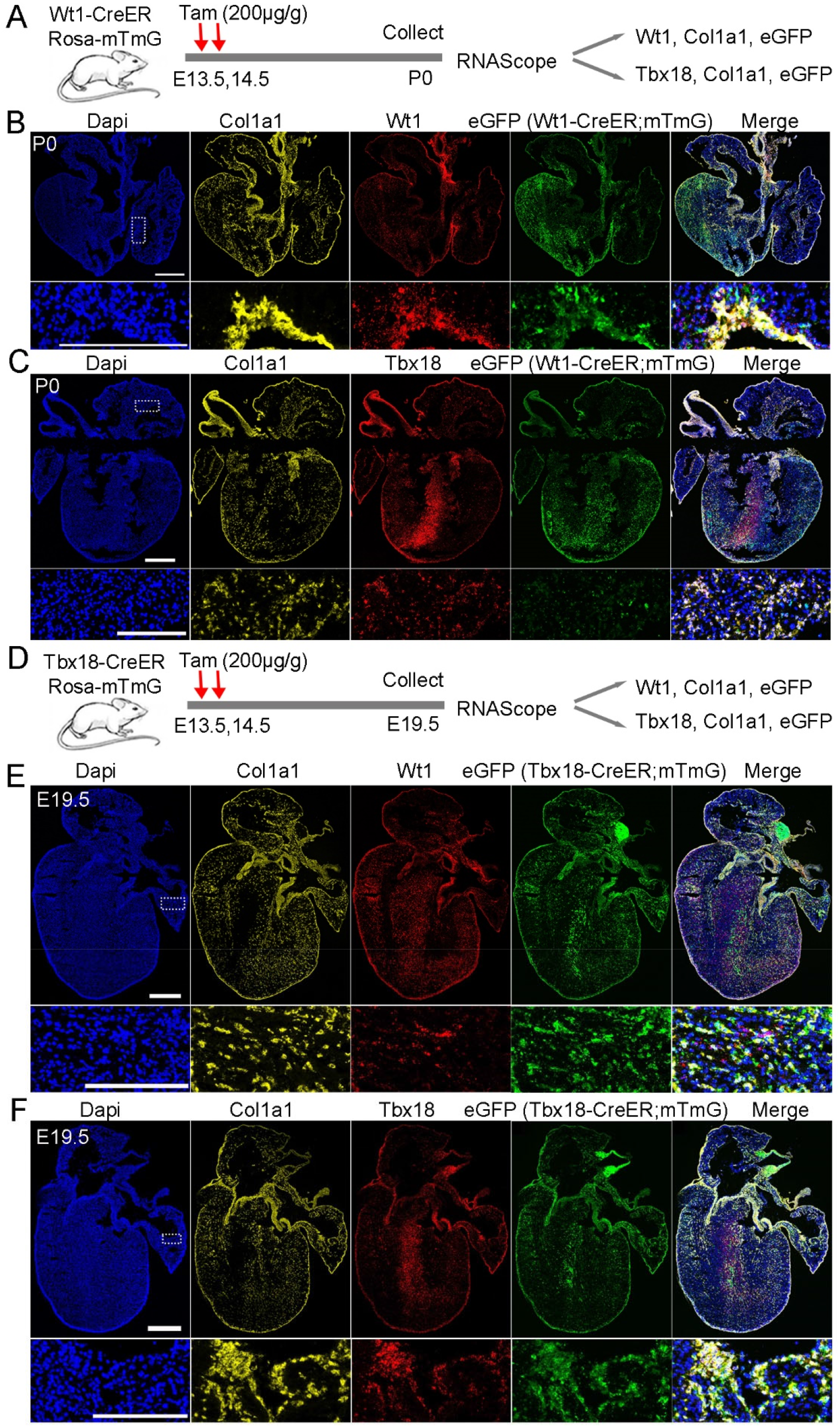
Lineage analysis of the Epi_FB subpopulations. (A) Diagram of the experimental workflow for staining the lineage descendants from Wt1-CreER;mTmG labeled cells. (B, C) RNAscope analysis of *Wt1* or *Tbx18*, *Col1a1*, and *eGFP* in fibroblasts derived from Wt1-CreER;mTmG labeled epicardial cells. (D) Diagram of the experimental workflow for staining the lineage descendants from Tbx18-CreER;mTmG labeled cells. (E, F) RNAscope analysis of *Wt1* or *Tbx18*, *Col1a1*, and *eGFP* in fibroblasts derived from Tbx18-CreER;mTmG labeled epicardial cells. Scale bar=500µm and 250µm in the whole heart images and enlarged areas, respectively.

### Comparative analysis of Postn-CreER and Pdgfra-CreER labeled lineages

To identify the right transgenic mice to label the entire cardiac fibroblast population, we analyzed the expression of two well-known fibroblast marker genes, *Postn* and *Pdgfra*, in the scRNA-seq data. We observed that *Postn* was highly expressed in fibroblasts and mural cells. Among fibroblasts, *Postn* was expressed in all subpopulations (Fig 5Ai). On the other hand, Pdgfra was specifically expressed in fibroblasts, although its expression was relatively lower than *Postn*, especially in postnatal staged atrial cells (Fig 5Aii, S6A, B). To evaluate their efficiency in labeling fibroblasts, we bred Postn-CreER mice with Rosa-mTmG reporter mice and administered tamoxifen (200µg/g) to pregnant mice at E13.5 (Fig 5Bi). We collected the hearts at E17.5 and observed only a small proportion of GFP-positive cells in the valves and septum (Fig 5Bii), suggesting a low labeling efficiency from the Postn-CreER mice. In contrast, we conducted the same experiments using Pdgfra-CreER mice and observed a significant proportion of GFP-positive cells in both the valves and heart chambers (Fig S6C, D). It is important to note that eGFP was detected through antibody staining, as the endogenous eGFP fluorescence was quenched during a specific procedure. Furthermore, we treated pregnant Pdgfra-CreER; Rosa-mTmG mice with two relatively low doses of tamoxifen (120µg/g) at E13.5 and E14.5, which resulted in a high proportion of GFP-labeled cells (Fig 5Ci, ii). These results demonstrate the efficient labeling capability of the Pdgfra-CreER; Rosa-mTmG mice.

**Fig 5.**
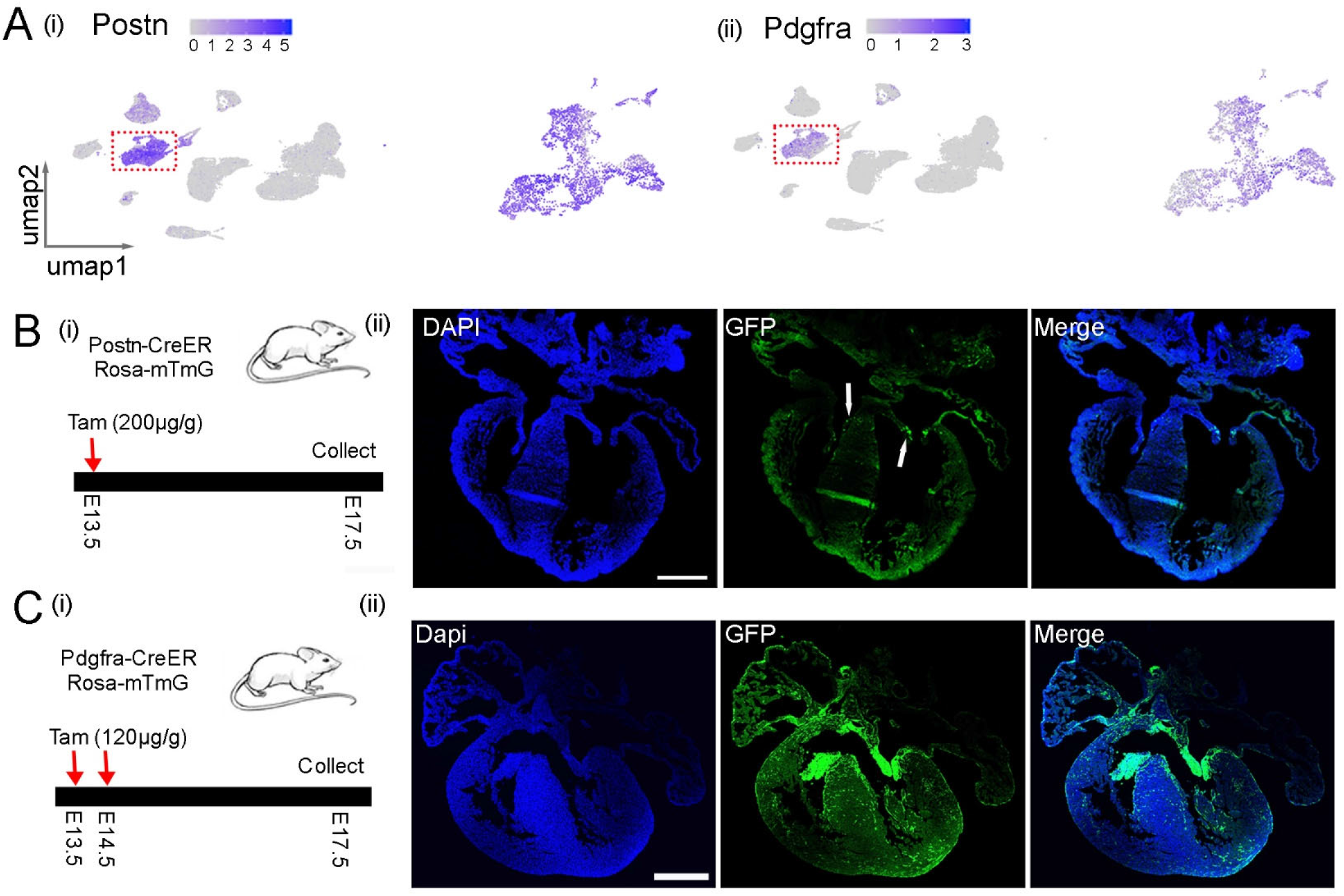
Lineage analysis of Postn-CreER and Pdgfra-CreER labeled cells. (Ai, ii) UMAP plots of *Postn* and *Pdgfra* expression in cardiac cells and cardiac fibroblasts. (B) (i) The workflow to lineage trace the Postn-CreER;mTmG labeled cells. (ii) A few cells in valves and septum were labeled in Postn-CreER; Rosa26-mTmG mouse hearts. (C) (i) The workflow to lineage trace the Pdgfra-CreER;mTmG labeled cells. (ii) An abundant of GFP+ cells were identified in the Pdgfra-CreER; Rosa-mTmG labeled mouse hearts. Scale bar = 500µm.

### Functional analysis of fibroblasts at different stages using a cell ablation system

Considering that cardiac fibroblast development was grouped into four phases, we were intrigued to understand its function in each of them. To do that, we utilized a genetically encoded DTA system, which involves the encoding of a toxin gene that induces apoptosis and allows for the ablation of target cells. Initially, we crossed floxed Rosa-DTA mice with Pdgfra-CreER mice and administered tamoxifen to pregnant female mice at E10.5. After 3 days, we harvested the embryos and observed that all the embryos with ablated fibroblasts (Pdgfra-CreER+/-; Rosa-DTA+/-) had perished, while the control embryos remained unaffected. This outcome indicates an essential role for fibroblasts in early-stage embryo development.

Subsequently, we administered tamoxifen to the female mice from the breeding pair at E13.5 and collected the embryonic hearts at E16.5 (Fig 6A). Our findings revealed that the ablated embryos exhibited smaller sizes compared to the control embryos, although the size of the ablated hearts did not significantly differ from that of the control hearts (Fig 6B). Additionally, we conducted TUNEL analysis to assess cell death in both control and ablated hearts and observed a significantly higher number of TUNEL-positive dots in the valve and chamber regions of the ablated hearts (Pdgfra-CreER+/-; Rosa-DTA+/-) compared to the control hearts, confirming the successful ablation of fibroblasts in the experimental hearts (Fig 6C, E). Furthermore, we examined the anatomical structure of the hearts but did not identify obvious defects in the ablated groups compared to control groups (Fig 6D). Additionally, we measured the thickness of the compact and trabecular myocardium in both the left and right ventricles. Our observations revealed a reduction in the thickness of the left ventricular (LV) compact myocardium and an increase in the thickness of the LV trabecular myocardium in the ablated hearts compared to the control hearts. Furthermore, we noted an elevated ratio of LV trabecular to compact myocardium in the ablated hearts compared to the control hearts (Fig 6F). However, no similar differences were observed in the right ventricle (RV). Finally, we examined cell proliferation post-ablation using pHH3 staining. We observed a reduction in pHH3-positive cells in the ventricular region, but not in the atrial region, of the ablated hearts compared to the control hearts (Fig 6G). These results suggest that cell proliferation in ventricle was impaired following fibroblast ablation.

**Fig 6.**
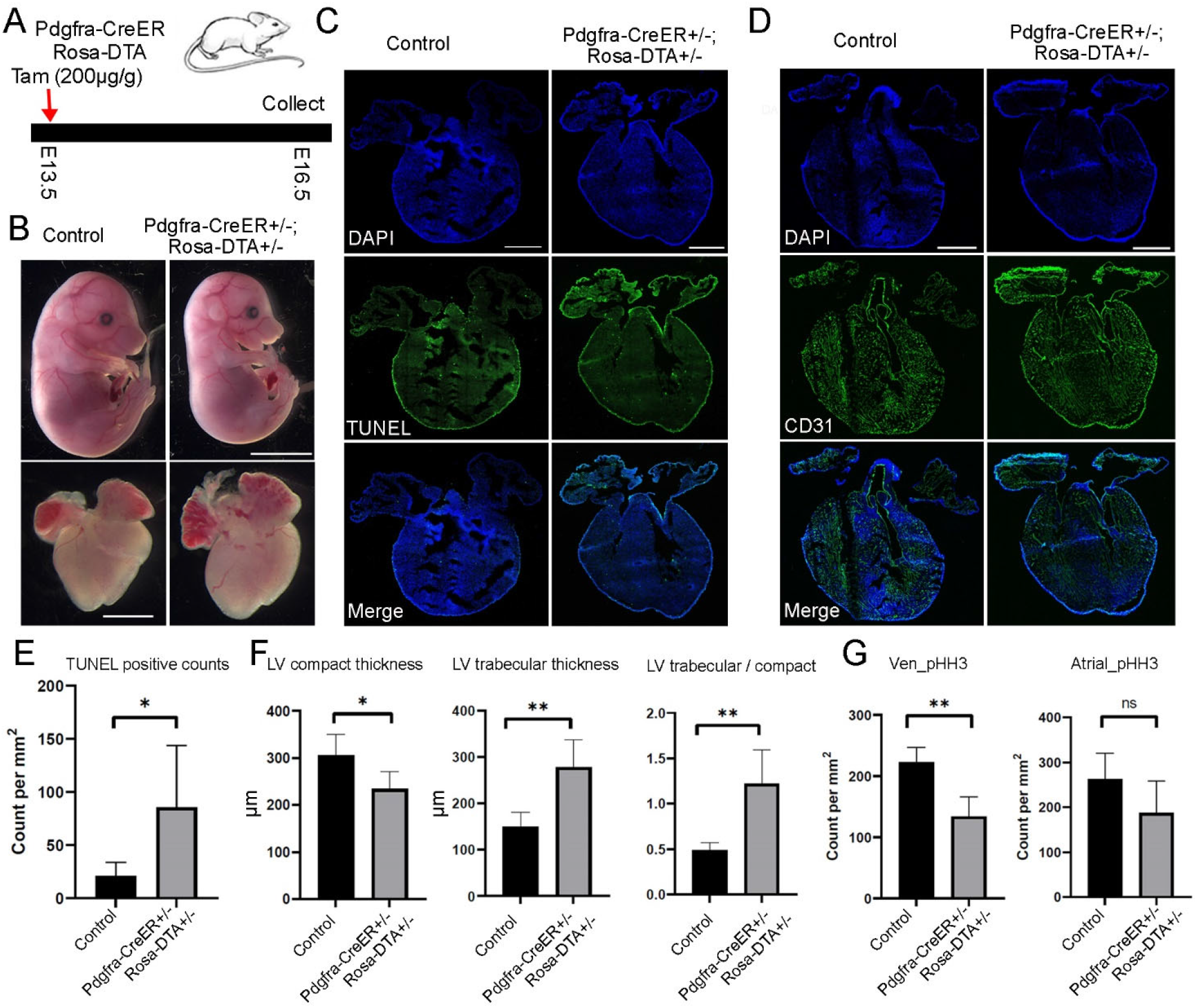
Analysis of fibroblast function with cell ablation at E13.5. (A) Diagram of the experimental procedure. (B) The ablated embryos were smaller than control embryos, but the ablated hearts were not obviously different from the control hearts. Scale bar=1mm. (C) Representative TUNEL staining results in control and ablated hearts. (D) Representative CD31 staining images in control and ablated hearts. (E) Quantification of TUNEL signal in control and ablated hearts. (F) Quantification of compact and trabecular myocardium thickness in control and ablated hearts. (G) Quantification of pHH3 signal in control and ablated heart ventricular and atrial regions. Scale bar =500µm. * represents p<0.05; ** represents p<0.01.

Next, we performed ablation in the third developmental phase by treating the pregnant mice with tamoxifen at E15.5 and harvested the embryos at E18.5 (Fig 7A). We did not observe obvious differences in the size of embryos and hearts between the ablated group and the control group (Fig 7B). However, the CD31 staining of heart sections revealed large holes in the ablated atrium but not in the ablated ventricles or control hearts (Fig 7C). Additionally, quantification of the compact and trabecular myocardium thickness in the LV and RV revealed a significant reduction in LV compact myocardium thickness but not in the other regions (Fig S7A). We also calculated the ratio of trabecular to compact myocardium thickness in the LV and RV and found that the ratio was increased in the RV but not in the LV in the ablated hearts compared to control hearts (Fig S7B). These results indicated that both the LV and RV in the ablated hearts have subtle defects. Moreover, we analyzed cell proliferation by staining pHH3 and found no significant differences between ablated and control ventricles (Fig S7C).

**Fig 7.**
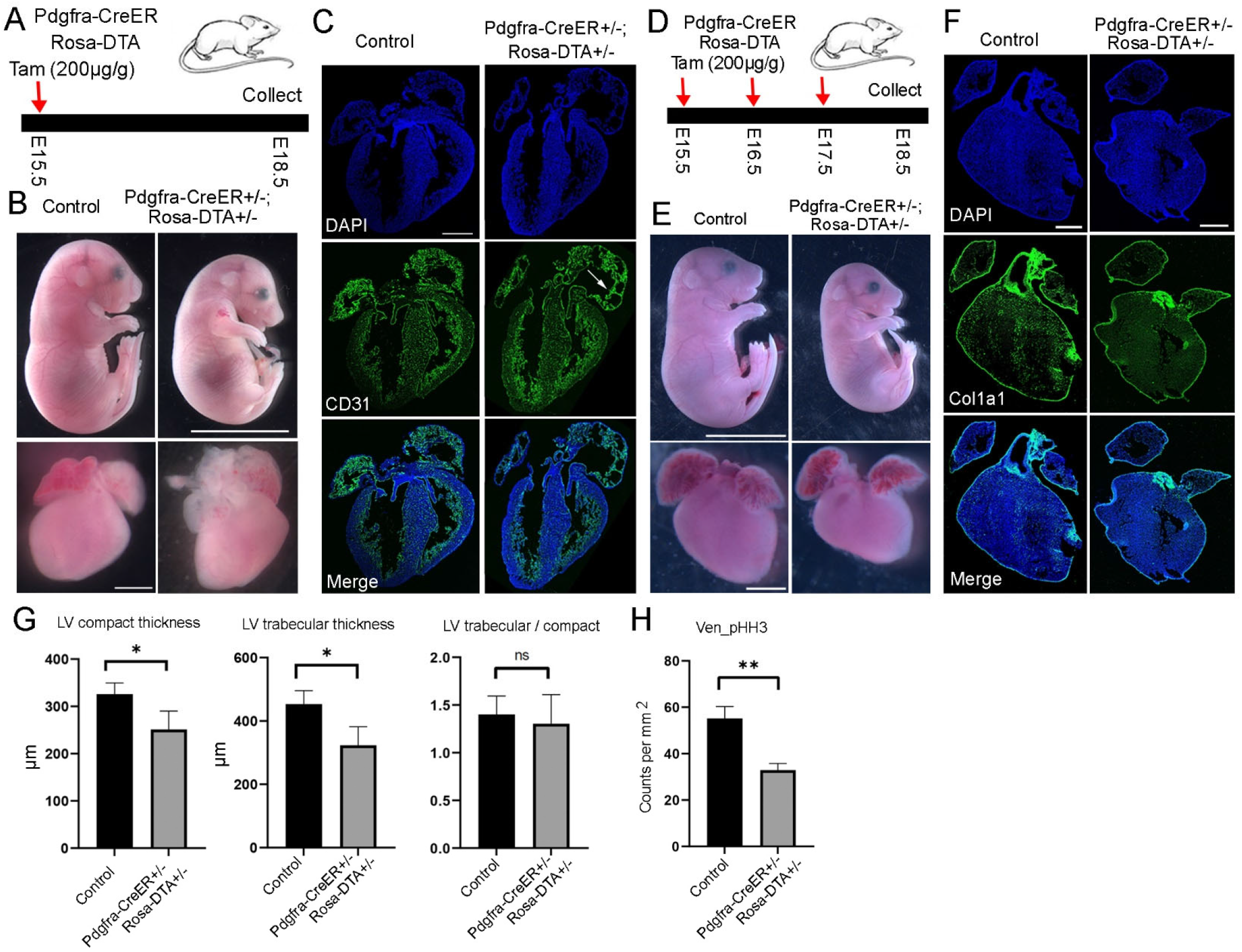
Functional analysis of fibroblasts at late embryonic stages. (A) Diagram of the experimental procedure with one dose of tamoxifen being administered at E15.5. (B) No obvious size differences were observed in the ablated embryos and hearts compared to controls. (C) CD31 staining analysis of control and ablated hearts. (D) Diagram of the experimental procedure with tamoxifen treatment from E15.5 to E17.5. (E) The ablated mice and hearts were smaller than controls. (F) RNA staining analysis of *Col1a1* in control and ablated hearts. The *Col1a1* signal was dramatically reduced in the ablated hearts. (G) Quantification of compact and trabecular myocardium thicknesses in control and ablated hearts. (H) Quantification of pHH3 signal in control and ablated hearts. Scale bar=1cm for the animal images and 500µm for the heart sections. * represents p<0.05; ** represents p<0.01.

Considering that the animals did not show obvious morphological changes after one dose of tamoxifen treatment, we increased the tamoxifen treatment to three consecutive days, ranging from E15.5 to E17.5 (Fig 7D). We harvested the embryos at E18.5 and found that the ablated embryos and their hearts were clearly smaller than the controls (Fig 7E). Next, to confirm the ablation efficiency, we analyzed *Col1a1* expression with RNA staining and found that the ablated hearts lost almost all of their Col1a1 signal in the four chambers (Fig 7F), indicating a high ablation efficiency with three doses of tamoxifen treatments. Further quantification of the LV compact and trabecular myocardium thickness showed a significant reduction in the ablated hearts compared to control hearts (Fig 7G). However, interestingly, the ratio of LV trabecular to compact myocardium thickness did not show differences between control and ablated hearts, indicating a proportional reduction of compact and trabecular myocardium thickness in the ablated hearts. Finally, we found that the ablated ventricles had less pHH3 signal than control ventricles (Fig 7H), suggesting that cell proliferation in the ventricles was impaired by fibroblast ablation.

Lastly, we assessed the fibroblast function in the fourth phase by treating the mice with tamoxifen at P1 and harvesting them at P4 (Fig 8A). We did not observe obvious differences between the ablated and control hearts (Fig 8B). However, CD31 staining of heart sections revealed defects in the atrium, which had a thinner and segmented atrial wall (Fig 8C). We further quantified the compact and trabecular myocardium in both the LV and RV but did not observe differences between control and ablated hearts (Fig S8A). We also did not find differences in the ratio of trabecular to compact myocardium thickness between control and ablated hearts in the LV and RV (Fig S8B). Finally, we analyzed cell proliferation by staining pHH3 and did not observe differences between the two conditions (Fig S8C). Next, we increased the tamoxifen treatments to three times, ranging from P1 to P3, and collected hearts at P4 (Fig 8D). The heart morphology and size did not reveal obvious differences between control and ablated groups (Fig 8E). The CD31 staining also did not show clear defects in the ablated hearts (Fig 8F). Finally, we quantified the chamber thickness and cell proliferation. Similar to hearts treated with one dose of tamoxifen, we observed no apparent defects in ablated hearts with three days of tamoxifen treatments (Fig 8G, S8D). In summary, the phase-specific ablation results revealed that fibroblasts played an essential role in early embryonic development, an important role in embryo and heart growth during the late embryonic stage, and a non-significant role in neonatal heart growth.

**Fig 8.**
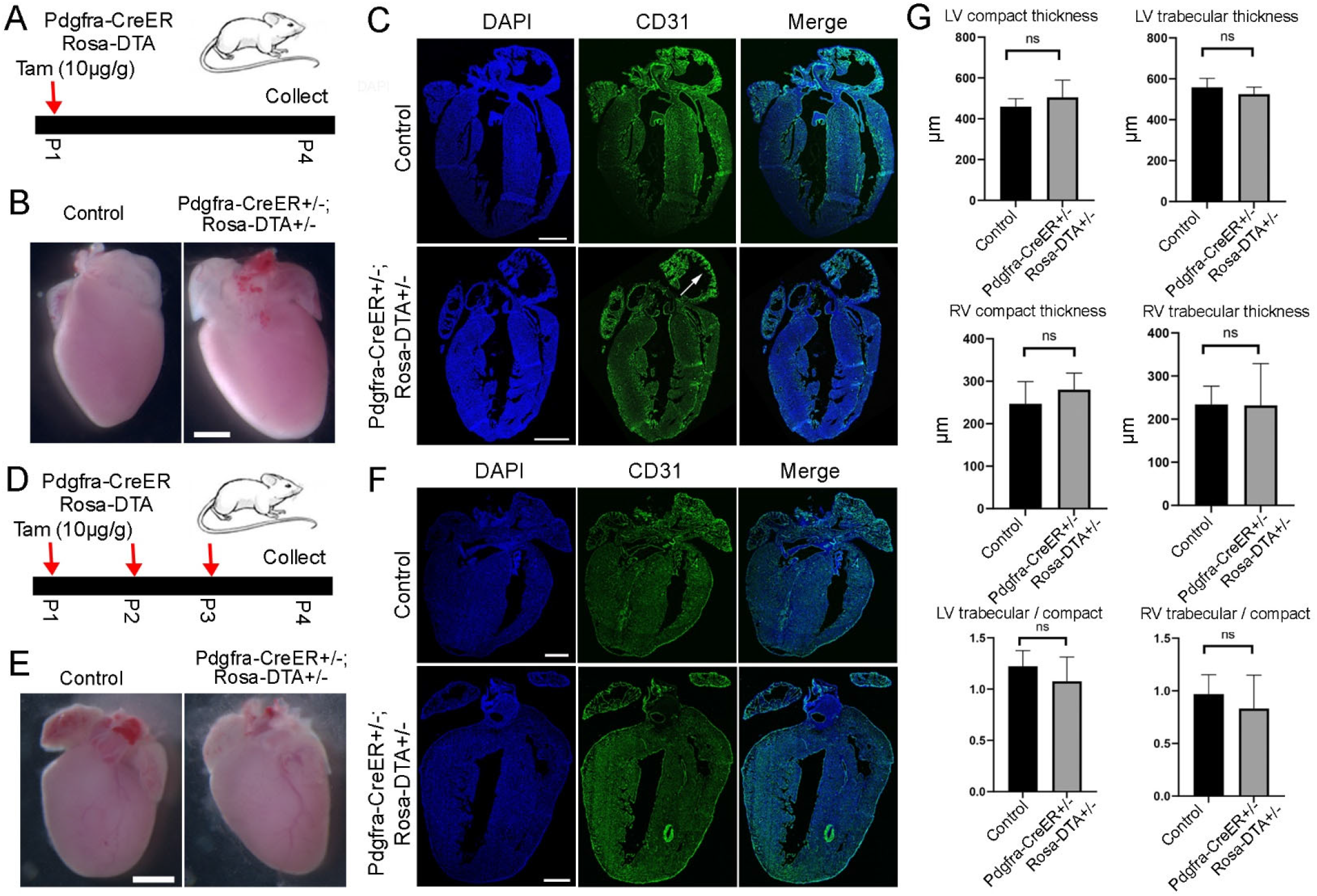
Functional analysis of fibroblasts at neonatal stage. (A) Diagram of the experimental procedure with tamoxifen treatment at P1. (B) No obvious size differences were observed in the ablated hearts compared to control hearts after one dose of tamoxifen treatment. (C) Representative images of control and ablated hearts stained with CD31. The defects in the right atrium from ablated hearts were pointed out by an arrow. (D) Diagram of the experiment to ablate fibroblasts with three doses of tamoxifen treatment from P1 to P3. (E) No obvious size difference was observed between control and ablated hearts. (F) Representative images of control and ablated hearts stained with CD31. (G) Quantification of the compact myocardium, trabecular myocardium, and their ratio in control and ablated hearts. Scale bar=500µm.

### Analysis of extracellular matrix genes expression and collagen accumulation along the developmental progression

Considering that one major function of fibroblasts is to secrete extracellular matrix proteins, we analyzed the expression pattern of the group of extracellular matrix genes (GO number, 0031012) in the main fibroblast population from CD1 and C57BL/6 mice. After calculating the enrichment score of each gene in each chamber and stage, we analyzed them with unsupervised clustering and identified five groups of genes that displayed stage or chamber specificities in both CD1 and C57BL/6 datasets (Fig 9A, S10A, Supplementary table S6, 7). Specifically, we found one group of genes (G1) that was highly expressed at stages before E14.5. This gene group includes signaling molecules such as *Slit2* and *Wnt5a*, and collagen genes such as *Col13a1* and *Col26a1* (Fig 9B, S10B). We also identified a group of genes (G2) that were mainly expressed in the LV and RV, which started to express at E14.5 and significantly increased its expression at the neonatal stage. This gene group includes collagen genes such as *Col5a1*, *Col6a1*, and *Col6a2*, and metalloproteinase genes such as *Adamts13* and *Adamts14* (Fig 9C, S10B). Additionally, we identified a group of genes that were highly expressed in the LA and RA at most stages, which includes *Col12a1*, *Adamts1*, and *Adamts4*. Interestingly, *Fn1* also exists in this group, suggesting that it has an important function in atrial fibroblast development (Fig 9D, S10B). Furthermore, we found a group of genes that were preferentially expressed in the other three chambers than the RA and were mainly expressed at late neonatal stages after P5 in certain chambers. This group of genes includes *Col4a4*, *Col4a5*, *Col4a6*, and *Ntn4* (Fig S9B, S10B). Finally, we identified a group of genes that displayed universal expression in all four chambers at most stages. This gene group includes *Col1a1*, *Col1a2*, and *Postn* (Fig 9B, S10B), which are the ubiquitously expressed extracellular matrix genes. In summary, we identified stage- and chamber-specific ECM genes in cardiac fibroblasts and observed a higher number of ECM genes with increased expression levels in the ventricular chambers during later stages.

**Fig 9.**
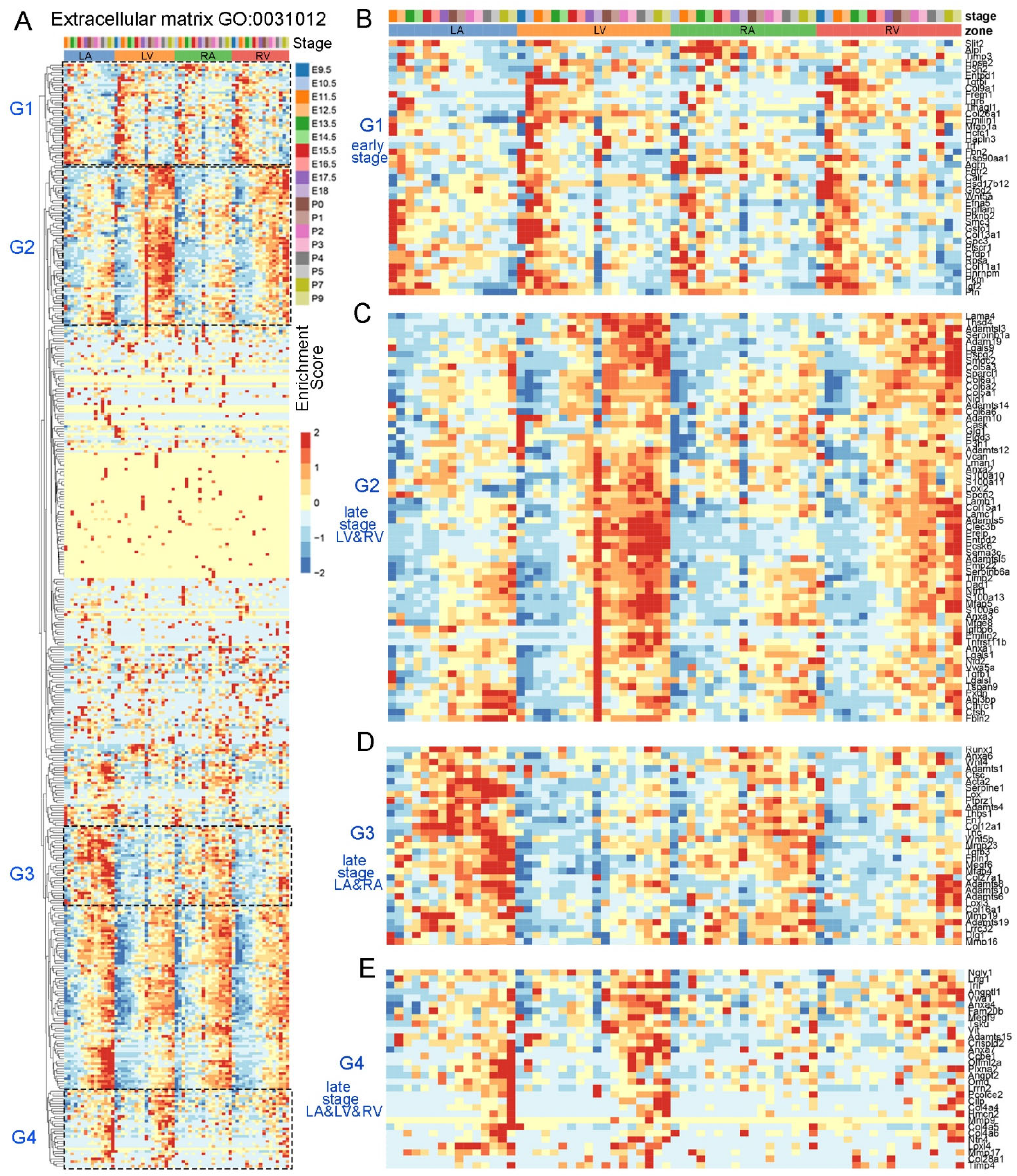
The expression pattern of extracellular matrix genes in the main population of cardiac fibroblasts. (A) Clustering analysis of the ECM gene expression enrichments. (B) The group of genes (G1) that were highly expressed at early staged fibroblasts in all four zones. (C) The group of genes (G2) that were highly expressed in LV and RV at late embryonic and neonatal stages. (D) The genes (G3) that were preferentially expressed in LA and RA. (E) The genes (G4) that were highly expressed in LA, LV, and RV at neonatal stage.

Given that collagen is one of the major ECM components and collagen genes are expressed at different stages and chambers, we analyzed collagen accumulation along the developmental progression. We did this by staining the tissue sections with collagen hybridizing peptide (CHP) after denaturing all collagens via an antigen retrieval process. Consistent with *Col1a1* expression, we observed strong collagen signals in the epicardium at all the analyzed stages (Fig S11A-D). Interestingly, at E11.5, we observed collagen within chambers besides the AVC (Fig S11A). Since fibroblasts in chambers have not developed yet at this stage, the collagen is likely derived from other cell types such as endocardial endothelial cells. At E14.5, we observed a strong collagen signal in the ventricular free walls and found the signal in the ventricular septum and atrium to be relatively weaker (Fig S11B). At E16.5, we found the collagen signal becoming brighter and denser in the entire heart, including the ventricular free wall, septum, and atrium (Fig S11C). This distribution was slightly broader compared to the fibroblast distribution at the same stage (Fig 1Aiii, iv). At P3, we found that the signal was further increased and formed stripe patterns (Fig S11D). In summary, we found that the progression of collagen accumulation largely coincided with the development of fibroblasts.

Next, we analyzed collagen in the hearts with fibroblast ablation. Firstly, we examined hearts at E18.5 (Pdgfra-CreER;Rosa-DTA mice and their littermate controls) treated with three times of tamoxifen from E15.5 to E17.5 (Fig S12A, B). We found that the collagen signal in the ablated hearts was sparser compared to the control hearts. Considering that most fibroblasts had been ablated in those hearts (Fig 7F), the remaining collagen signal was either derived from other cell types such as endothelial cells or generated by fibroblasts before ablation. Furthermore, we stained the P4 hearts with tamoxifen treatments from P1 to P3 (Fig S12C, D). We observed that the ablated hearts had a sparser collagen signal and lost most of the stripe pattern in the septum compared to the control hearts. However, a significant amount of stripe-patterned collagen was still observed in the ventricular free wall, although the overall signal was sparser. These collagens were likely derived from fibroblasts before ablation and could contribute to the insignificant heart defects after fibroblast ablation at this phase.

### Identification of stage enriched ligand-receptor interactions between fibroblast and cardiomyocyte

Fibroblasts can also contribute to heart development by interacting with other cell lineages. To study this process, we conducted an analysis of ligand-receptor interactions between fibroblasts and atrial or ventricular CMs. We analyzed the interactions at each stage and identified those that were unique to a particular stage, as well as those that were shared by multiple stages (Supplementary table S8, 9). Interestingly, for the stage-unique interactions between Fb and Ven_CM, we found that they were predominantly present during early embryonic stages and the neonatal stage. We did not find any unique interactions between E13.5 and E17.5 (Fig S13). However, for the unique interactions between Fb and Atrial_CM, we observed relatively even numbers across stages (Fig S14). Additionally, we observed that the interactions in the atrium and ventricle at each stage largely did not overlap. These results suggest that fibroblasts in the atrium and ventricle have distinct signaling communications with CMs.

Next, we analyzed the interactions that are shared by multiple stages. We found that most of them were enriched at early embryonic stages, ranging from E9.5 to E14.5, in both atrial and ventricular CMs (Fig 10A, B). We observed a few interactions between E15.5 to E17.5, which were mostly also expressed at the early or neonatal stage. At the neonatal stage, we noticed a relatively higher number of interactions. Overall, we observed the highest number of interactions at early embryonic stages before E15.5, the least interactions at late embryonic stages after that, and a moderate number of interactions at the neonatal stage. This aligns well with cardiac lineage specifications during early stages and the adaptation to environmental changes associated with birth during the neonatal stages. Additionally, we identified more interactions in the ventricle compared to the atrium. Moreover, most of these interactions were not shared between the two regions (Fig 10A, B), further confirming the existence of distinct signaling interactions between FBs and CMs in these two areas.

**Fig 10.**
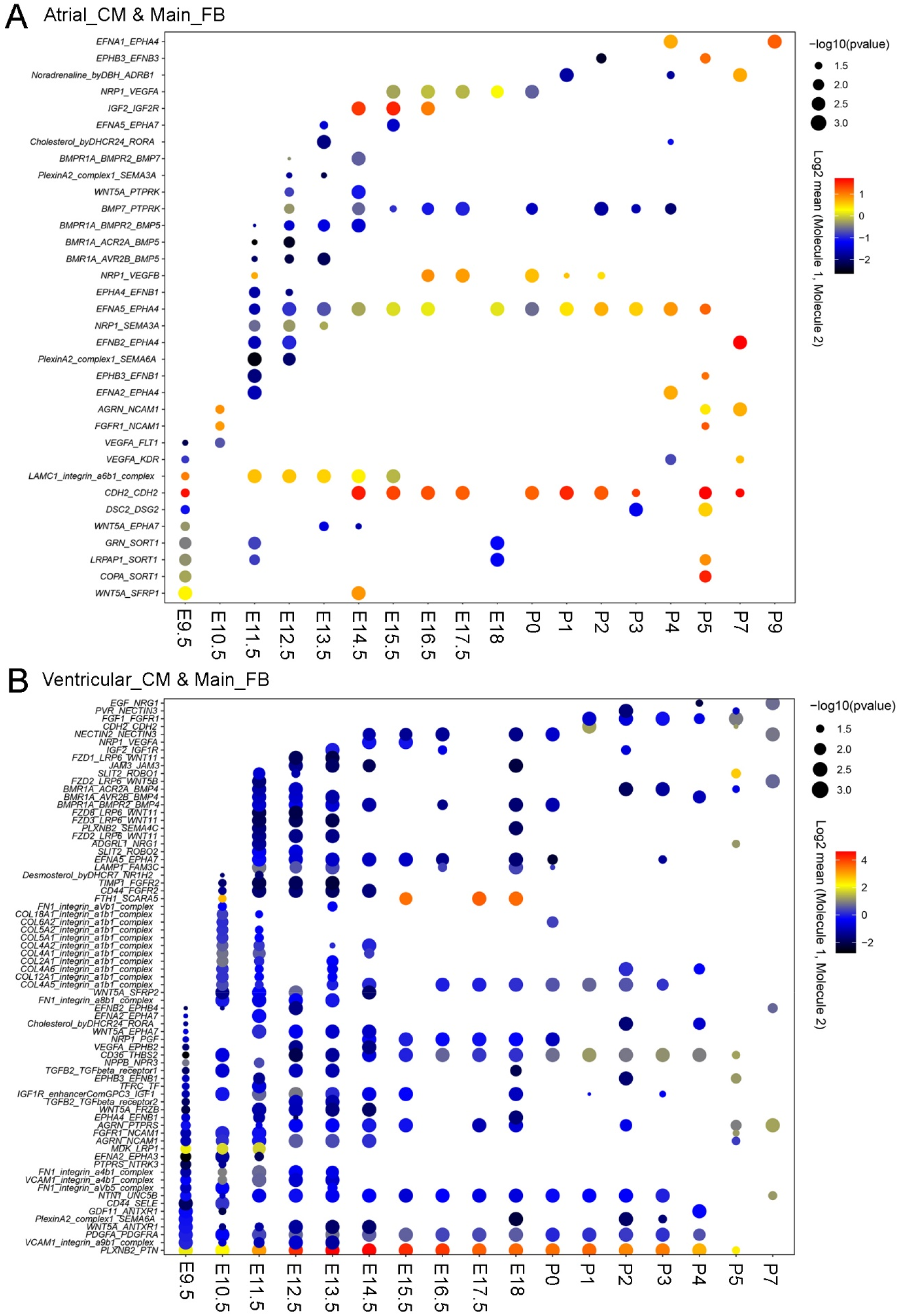
The ligand-receptor interactions between FBs and atrial or ventricular CMs that expressed at multiple stages. (A, B) The interactions that are shared by at least two stages between the main population of fibroblasts and atrial or ventricular CMs.

## Discussion

In this study, we identified heterogeneity in cardiac fibroblasts and found that the subpopulations largely preserved the expression of lineage genes likely from their precursor cells. We also grouped cardiac fibroblasts into different developmental phases based on their developmental progression, and further studied their function at each phase using a cell ablation system. Finally, we identified the extracellular matrix genes and ligand-receptor interactions that were expressed at each stage and chamber. Altogether, our study demonstrates the first comprehensive study of fibroblast heterogeneity and function in heart development.

In our scRNA-seq data, we only identified a small proportion of *Cdh5*-positive fibroblasts. Further RNA staining analysis showed that they are located at the root of large vessels. However, according to previous lineage tracing results^9^, the endocardial endothelial cell lineage contributes to most fibroblasts in the ventricular septum. This suggests that most fibroblasts in the septum have lost their endothelial cell gene expression signature at the neonatal stage. Next, it would be interesting to analyze if the cluster of *Cdh5*-positive fibroblasts that we have identified in the study will also downregulate the endothelial cell gene expression at later stages. Additionally, we found that *Sox10*-positive fibroblasts are also located around corollary vessels, in addition to large vessels. It would be interesting to analyze their functions in corollary vessel development and physiology in the future.

The atrial fibroblasts developed relatively late and were mainly derived from *Wt1*- and *Tbx18*-positive epicardial cells. RNA staining showed that *Wt1*-positive fibroblasts were mainly located on the edge, while *Tbx18*-positive fibroblasts filled the whole chambers. It would be interesting to know where the two types of epicardial cells are located in the atrial epicardium and when they become distinct from each other. Additionally, the scRNA-seq data showed that *Wt1*-positive fibroblasts in the atrium were mainly in the G1 phase, suggesting that they are less proliferative. Considering that these cells are also physically adjacent to the atrial epicardium, they are likely newly formed cells from epicardial cells via epithelial-mesenchymal transition (EMT). It would be interesting to understand their function and if they will become active in cell proliferation later.

The scRNA-seq data showed that *Postn* was more highly expressed in fibroblasts than *Pdgfra*, but the lineage tracing results revealed a lower labeling efficiency. Inconsistency between gene expression and labeling efficiency is not uncommon in CreER mice driven by gene loci. Additionally, our scRNA-seq data (Fig S6A, B) showed that *Pdgfra* expression was reduced in a subset of atrial fibroblasts at the neonatal stage. However, it is unknown if this reduction would cause a reduction in labeling efficiency in those cells. Moreover, most fibroblast genes like *Pdgfra* and *Tcf21* are expressed in multiple organs. This makes cell or gene ablation using the transgenic mice Pdgfra-CreER and Tcf21-CreER not tissue-specific. To specifically label and ablate fibroblasts in the heart, we have at least two options. The first option is to identify genes that are uniquely expressed in cardiac fibroblasts and then generate CreER mouse lines based on them. Another option is to develop a dual recombination system with one recombinase, such as Cre, that expresses specifically in fibroblasts and another recombinase, such as Dre, that expresses specifically in the heart^28, 29^. The combination of the two recombinases can achieve heart and fibroblast specificity. Once the system is developed, it also has the potential to label the cardiac fibroblast subpopulations that we have identified in the study.

Cell ablation systems have been widely used to study tissue regeneration in different species. In the heart, DTA-based cell ablation has been used to ablate cardiac progenitors and cardiomyocytes, and has revealed a robust regeneration response after cell ablation at early stages^22^. However, in our study, based on the pHH3 staining signal, we did not observe an increase in cell proliferation after cell ablation. This could be due to multiple possibilities. First, the Pdgfra-CreER; Rosa-DTA-based ablation may be too severe to initiate the regeneration response. Considering that ablation at E13.5 and ablation with three doses of tamoxifen at E15.5 both led to smaller embryos, the ablation of a large number of cells may compromise the regeneration/compensation mechanisms. However, it is difficult to explain the embryos at E15.5 with one dose of tamoxifen treatment, which did not display obvious defects after ablation and have no increase in cell proliferation. Second, the ablation stages may be relatively late, and fibroblasts may have lost their regeneration capability. Third, fibroblasts may be different from other cardiac cells and have a limited regeneration response to their own ablation, although they can respond to cardiomyocyte loss at neonatal and adult stages.

Through ablation at each phase, we found that FBs play crucial roles in early embryo survival and late embryo growth, but their significance diminishes in neonatal heart growth. This could be partially explained by our discovery of a large number of ligand-receptor interactions at early stages and the high levels of ECM gene expression and collagen accumulation observed during the neonatal stage. Furthermore, since neonatal hearts mainly develop through CM hypertrophy and maturation rather than cell proliferation, fibroblasts may primarily function to promote these aspects. Additionally, it’s worth noting that our analysis was conducted shortly after ablation, which allowed us to observe defects during fast growth periods like embryonic stages, but may not have been sufficient to capture relatively slower growth phases like the neonatal stage. Moreover, we noticed that the atrium appears to be more susceptible to defects after ablation compared to the ventricle during later embryonic and neonatal stages. This observation can likely be explained by the relatively delayed development of the atrium compared to the ventricle.

One critical role of fibroblasts is to secrete extracellular matrix proteins. The scRNA-seq data showed that there are many types of ECM proteins besides collagens, and further work will be required to analyze their temporal and spatial accumulation and function in heart development. Additionally, we found that most ECM genes in group 4 upregulate their expression at postnatal day 7, when the mouse hearts start to lose their regeneration capability^30^. Next, it will be interesting to test their function in heart regeneration by inhibiting their expression at that time point. Furthermore, fibroblasts can also interact with other cell types to regulate heart development and homeostasis. In this study, we mainly focused on its interactions with cardiomyocytes, but it will also be interesting and important to study its interactions with the other cardiac cell types, such as endothelial cells and immune cells, in the future.

## Methods

### Experimental methods

#### Mouse strains

The animal experiments have been approved by the University of Pittsburgh Institutional Animal Care and Use Committee (IACUC). CD1 male and female mice were purchased from Charles River Laboratories and bred in our laboratory to generate embryos and neonatal pups at specific stages for RNA staining experiments. The transgenic mice, including Wt1-CreERT2 (Strain #:010912)^31^, Tbx18-CreERT2 (Strain #:031520)^32^, Rosa26-mTmG (Strain #:007676)^33^, Pdgfra-CreERT2 (Strain #:032770)^34^, Postn-CreER (Strain #:029645)^35^, and ROSA26-eGFP-DTA (Strain #:032087)^20^, were ordered from the Jackson Laboratory.

#### Tamoxifen treatment and mouse dissection

The male and female mice of specific strains were bred together. The default dosage of 200µg of tamoxifen per gram of body weight (200µg/g) was given to pregnant mice through oral gavage, and neonatal mice were given 10µg/g of tamoxifen by direct injection into their stomach to induce Cre activity^23^. The only exception is that 120µg/g of tamoxifen was given to the Pdgfra-Cre;Rosa-mTmG pregnant mice at E13.5 and E14.5 (Fig 5C). The pregnant mice and neonatal pups were euthanized using CO2 and decapitation-based methods, respectively. The mouse hearts were isolated following the standard procedure described previously and were directly embedded in OCT without fixation for RNA staining or fixed at 4% paraformaldehyde for other experiments, such as immunofluorescence staining.

#### Proximity ligation in situ hybridization (PLISH)

PLISH was performed following a published protocol^36^. Specifically, the embryonic or postnatal mouse hearts were embedded in OCT (Sakura, 4583) without fixation. After sectioning at a thickness of 10μm, the tissue sections were treated with post-fix medium (RNase-free PBS with 3.7% formaldehyde and 0.1% DEPC) followed by 0.1mg/ml pepsin treatment (RNase-free H2O with 0.1mg/ml pepsin and 0.1M HCl). After dehydration, the sections were sealed with hybridization chambers (Invitrogen, S24732) and hybridized with H probes (supplementary table S1) in Hybridization Buffer (1 M NaTCA, 5 mM EDTA, 50 mM Tris pH 7.4, 0.2 mg/mL Heparin). Next, after being treated with circularization reaction and rolling cycle amplification, the samples were hybridized with detection probes conjugated with Cy3 or Cy5 fluorophore. Finally, the samples were stained with DAPI (Invitrogen, D1306), mounted with fluoromount-g (SouthernBiotech, OB100-01), and imaged under confocal microscopy (Leica TSC SP8).

#### RNAscope Multiplex Fluorescent V2 Assay

The RNAscope Multiplex Fluorescent Reagent Kit v2 (Advanced Cell Diagnostics, 323270) was performed according to the manufacturer’s manual. Briefly, the tissue sections were fixed in pre-chilled 4% PFA (Electron Microscopy Sciences,15710) at 4℃ for 1 hour. After progressive dehydration, the sections were sequentially treated with hydrogen peroxide for 10 minutes and protease IV for 15 minutes (for embryonic) or 20 minutes (for postnatal) at room temperature (RT). After that, the samples were hybridized to gene-specific Z probes for 2 hours at 40℃ using the HybEZ II Hybridization system (ACD, 321721). Following further signal amplification, the hybridization signals were detected with TSA Vivid Fluorophores. The samples were then stained with DAPI and mounted with ProLong Gold Antifade Mountant (Invitrogen, P36930), and imaged under confocal microscopy (Leica TSC SP8). The RNAscope probes used in the study include Mm-Cdh5-C3 (312531-C3), Mm-Sox10-C3 (435931-C3), EGFP-O4 (538851), Mm-Col1a1-C2 (319371-C2), Mm-Wt1-C3 (432711-C3), and Mm-Tbx18-C3 (515221-C3).

#### Immunofluorescence staining

The immunofluorescence staining was performed following a standard procedure. Briefly, mouse hearts were fixed in 4% PFA overnight before embedding in OCT. Afterwards, the samples were sectioned at 10µm and briefly washed in PBS to clean the OCT. The sections were then blocked for 1 hour in blocking buffer (10% goat serum, 1% BSA, 0.1% Tween 20) and incubated with primary antibodies in the primary antibody buffer (1% BSA in PBST) at 4℃ overnight. On the second day, the samples were stained with fluorophore-conjugated secondary antibodies in blocking buffer for 1 hour at room temperature. Finally, the samples were stained with DAPI, mounted with fluoromount-g, and imaged with a confocal microscope. The primary antibodies used in the study include anti-eGFP (ThermoFisher, #CAB4211), anti-CD31 (BD, # 550274), and anti-pHH3-488 (abcam, #ab197502). The secondary antibodies used are Goat Anti-Rabbit IgG-488 (ThermoFisher, # A-11008) and Goat Anti-Rabbit IgG-647 (ThermoFisher, #A21247).

#### Collagen staining

Fresh frozen sections from embryonic and postnatal mouse hearts were fixed in 4% PFA in 1X PBS for 15 minutes at room temperature (RT). After that, the sections were soaked in a sodium citrate-based solution (distilled water with 10mM sodium citrate and 0.5% Tween-20, HCl was added to adjust pH to 6.0) for antigen retrieval, which was performed in a steamer for 30 minutes. Subsequently, the sections were incubated with 20 µM biotin-conjugated collagen hybridizing peptide (Advanced Biomatrix, 50-196-0307) in 1X PBS at 4℃ overnight. The next day, streptavidin-cy5 (Invitrogen, SA1011) was used (1:500 in 1X PBS with 1% BSA) for 1 hour at RT. Finally, the samples were stained with DAPI, mounted with FLUOROMOUNT-G, and imaged under a confocal microscope.

#### TUNEL staining

TUNEL staining was performed following the manufacturer’s protocol (Roche, 11684795910). Briefly, mouse hearts were embedded and sectioned as described in the immunofluorescence staining procedure. After fixation with 4% Paraformaldehyde in PBS for 20 minutes and three washes with PBS, the sections were incubated in permeabilization solution (0.1% Triton X-100, 0.1% sodium citrate) for 2 minutes on ice, followed by two PBS rinses. Subsequently, 60 µl of TUNEL reaction mixture was added to each section, and they were incubated in a humidified atmosphere for 1 hour at 37℃ in the dark. Finally, the sections were washed with PBS three times, stained with DAPI, mounted with fluoromount-g, and imaged with a confocal microscope.

#### Statistical analysis

To quantify pHH3 signal, compact, and trabecular myocardium thickness, we used two to three sections from each heart and two to four hearts per genotype. Statistical analyses were conducted using ImageJ and Prism 9 software, employing a two-tailed Student’s t-test to compare groups. P-values below 0.05 were considered significant.

### Data analysis

#### Data Availability

The scRNA-seq datasets from CD1 and C57BL/6 mouse hearts were generated in the previous study^26^ and can be downloaded from the Gene Expression Omnibus (GEO) database using the accession number GSE193346.

#### Fibroblast scRNA-seq data subset, clustering, and gene expression analysis

We mainly used Seurat V4 for this part of the analysis^37^. Firstly, we directly plotted the expression of genes including Col1a1, Pdgfra, and Postn using the CD1 and C57BL/6 Seurat objects. Next, we selected the fibroblast-like cells from the objects using the “subset” function and further re-clustered them following the vignette on the Seurat website with default settings.

Once we had the Seurat objects with fibroblasts only, we plotted the gene expression using FeaturePlot and annotated their cell cycle phases using the CellCycleScoring function. We then identified the genes that were differentially expressed among the clusters using the “FindAllMarkers” function with default settings. Next, we repeated the same procedure to analyze the main fibroblast populations to understand their heterogeneities. To plot the cell numbers in each cluster, we used the “table(Idents(objects))” function in Seurat to identify the values and then generated the plots in Prism 9 (GraphPad Software, Inc).

To analyze the valve cells from CD1 and C57BL/6 datasets, we first extracted the gene expression profile of these cells from both datasets using the “subset” function. Next, we integrated them together using the “merge” function. After the standard procedure, including data normalization, scaling, and PCA analysis, we used the package Harmony^38^, following its standard procedure with default settings, to remove potential batch effects between mouse strains. Finally, we plotted the valve cells with different labels, including cluster, cell cycle phase, stage, and zone, using the Seurat function DimPlot. We also plotted the expression of representative genes using FeaturePlot.

#### Extracellular matrix genes expression analysis

We downloaded the extracellular matrix genes from the Jackson Laboratory under the term “extracellular matrix” and gene ontology ID 0031012 (https://www.informatics.jax.org/go/term/GO:0031012; Download date: 2023-07-07). After downloading, we cleaned the gene list by removing duplicates and filtering out genes with null expression, resulting in a total of 440 genes. Next, we assessed the enrichment of each gene in the main population of fibroblasts at each stage and chamber using the R package AUCell^39^, with heatmaps drawn with the R package ggplot2 (Fig 9, S9, S10).

#### Ligand-receptor interaction analysis

To identify ligand-receptor interactions across stages and zones for fibroblasts and cardiomyocytes, we analyzed relevant subsets of the CD1 data, including the main population of fibroblasts identified in figure 2D, atrial cardiomyocytes, and ventricular cardiomyocytes, using CellPhoneDB v3.34. We used a p-value threshold of 0.2 and ran the analysis with 10 threads, while keeping the rest of the parameters at their default settings. The significant mean value of all interactive partners (log2) and enrichment p-values (-log10) obtained from the CellPhoneDB outputs were plotted as dotplots in R.

## Acknowledgements

We’d like to thank all the members in the Li laboratory for their insightful discussions of this work. This research was supported in part by the University of Pittsburgh Center for Research Computing, RRID:SCR_022735, through the resources provided. Specifically, this work used the HTC cluster, which is supported by NIH award number S10OD028483.

## Author Contributions

Y.D., Y.H., G.L. designed the experiments; Y.D. analyzed the ablated hearts with different assays; J.X. bred the mice, treated them with tamoxifen, and harvested the hearts from embryonic and neonatal mice. Y.H. performed the RNAScope and collagen staining experiments, Y.H. and G.L. performed the PLISH experiments; H.T. and G.L. analyzed the scRNA-seq data; Y.D., Y.H., H.T., and G.L. prepared the manuscript; All the authors edited the manuscript.

## Legends for Supplemental Figures

Supplementary Fig S1: Identification of different cardiac fibroblast populations in C57BL/6 mice with scRNA-seq. (A) UMAP plots showing the expression of representative lineage genes. (B, C) UMAP plots of scRNA-seq data labeled by stage or cell cycle phases. (D) Summary of the cardiac fibroblast populations in C57BL/6 mice that we have identified with scRNA-seq.

Supplementary Fig S2: Integrative analysis of the valve interstitial cells from CD1 and C57BL/6 scRNA-seq datasets. (A-D) UMAP plots of the VICs labeled by cluster, cell cycle phases, stage, and zone. (E-G) UMAP plots showing the expression of representative VIC genes.

Supplementary Fig S3: Stage analysis of the cells in each main FB subpopulation. (Ai-v) The stage distribution of each subpopulation in G1 phased FBs (CD1 mice). (Bi, ii) The stage distribution of each subpopulation in S_G2M phased FBs (CD1 mice).

Supplementary Fig S4: The subpopulations in C57BL/6 main FBs. (A-C) UMAP plots of main population of FBs labeled by stage, zone, and cell cycle phases. (D) UMAP plots showing the expression of representative epicardial lineage genes *Wt1*, *Tbx18*, and *Aldh1a2*. (E) The subpopulations within C57BL/6 main FBs.

Supplementary Fig S5: Confirmation of FB subpopulations through RNA staining. (A) Co-staining of *Col1a1* and *Wt1* with RNAScope at E17.5. (B) Co-staining of *Col1a1* and *Tbx18* with RNAScope at E17.5. (C) Co-staining of *Col1a1* and *Wt1* with PLISH at E17.5. (D) Co-staining of *Col1a1* and *Tbx18* with PLISH at P3. Scale bar=500µm in the whole heart sections; Scale bar=250µm in the zoomed sections in A and B; Scale bar=100µm in the zoomed section in C and D.

Supplementary Fig S6: (A, B) UMAP plots of FBs in CD1 mice labeled by stage and zone. (C, D) Lineage analysis of Pdgfra-CreER+/-; mTmG+/-mouse hearts with 200ug/g of tamoxifen treatment at E13.5. Scale bar=500µm.

Supplementary Fig S7: Quantification of the defects in control and ablated hearts with one dose of tamoxifen treatment at E15.5. (A) Quantification of the compact and trabecular myocardium thickness in LV and RV. (B) Quantification of the ratio of trabecular to compact myocardium in LV and RV. (C) Quantification of pHH3 positive cells in control and ablated ventricles. * represents p<0.05; ** represents p<0.01.

Supplementary Fig S8: Quantification of the defects in control and ablated hearts at neonatal stage. (A) Quantification of the compact and trabecular myocardium thickness in LV and RV after one dose of tamoxifen treatment. (B) Quantification of the ratio of trabecular to compact myocardium in LV and RV after one dose of tamoxifen treatment. (C) Quantification of pHH3 positive cells in ventricular in control and ablated hearts after one dose of tamoxifen treatment. (D) Quantification of pHH3 positive cells in ventricular in control and ablated hearts after three doses of tamoxifen treatments. * represents p<0.05; ** represents p<0.01.

Supplementary Fig S9: (A, B) The expression pattern of a group of genes (G5) that displayed expression in all four chambers of FBs (CD1 mice) at late embryonic and neonatal stages.

Supplementary Fig S10: The expression pattern analysis of extracellular matrix genes in C57BL/6 FBs. (A) Unsupervised clustering analysis of extracellular matrix genes expression in C57BL/6 FBs. (B) The groups of genes that display stage or zone-specific expression pattern.

Supplementary Fig S11: Staining analysis of collagen accumulation in different stages of hearts using collagen hybridizing peptide. (A-D) Collagen expression pattern in CD1 mouse hearts at E11.5, E14.5, E16.5, and P3. Scale bar=500um and 150um in the whole heart sections and enlarged sections, respectively.

Supplementary Fig S12: In situ analysis of collagen accumulation in control and ablated hearts. (A) Collagen accumulation in control hearts at E18.5. (B) Collagen accumulation in Pdgfra-CreER;Rosa-DTA ablated hearts. Three doses of tamoxifen were given at E15.5, E16.5, and E17.5. (C) Collagen accumulation in control hearts at P4. (D) Collagen accumulation in Pdgfra-CreER;Rosa-DTA ablated hearts. Three doses of tamoxifen were given at P1, P2, and P3. Scale bar=500µm and 150µm in the whole heart sections and enlarged sections, respectively.

Supplementary Fig S13: The stage-unique ligand receptor interactions between the main population of fibroblast and ventricular CMs.

Supplementary Fig S14: The stage-unique ligand receptor interactions between the main population of fibroblast and atrial CMs.

## Legend for Supplementary Tables

Supplementary Table 1: Primer sequences for PLISH probes

Supplementary Table 2: The differentially expressed genes among fibroblast clusters in CD1 dataset.

Supplementary Table 3: The differentially expressed genes among fibroblast clusters in C57BL/6 dataset.

Supplementary Table 4: The differentially expressed genes among clusters in CD1 main population dataset.

Supplementary Table 5: The differentially expressed genes among clusters in C57BL/6 main population dataset.

Supplementary Table 6: The enrichment score of extracellular matrix genes in CD1 main population fibroblasts.

Supplementary Table 7: The enrichment score of extracellular matrix genes in C57BL/6 main population fibroblasts.

Supplementary Table 8: The list of ligand-receptor interactions between the main population of fibroblast and atrial cardiomycocytes in CD1 dataset.

Supplementary Table 9: The list of ligand-receptor interactions between the main population of fibroblast and ventricular cardiomycocytes in CD1 dataset.

