## Supplemental Figures for "Heterogeneity and Functional Analysis of Cardiac Fibroblasts in Heart Development"

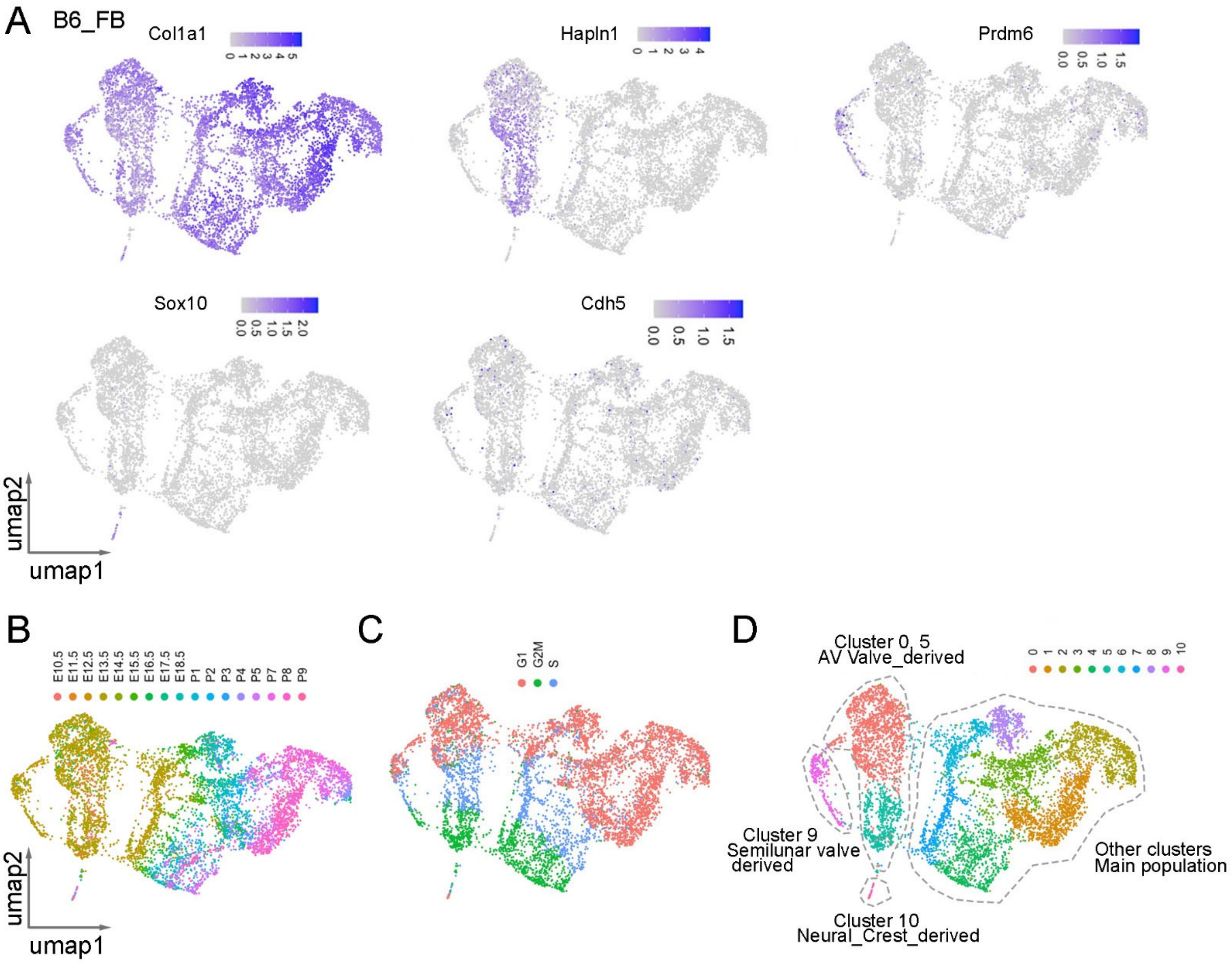


Supplementary Fig S1: Identification of different cardiac fibroblast populations in C57BL/6 mice with scRNA-seq. (A) UMAP plots showing the expression of representative lineage genes. (B, C) UMAP plots of scRNA-seq data labeled by stage or cell cycle phases. (D) Summary of the cardiac fibroblast populations in C57BL/6 mice that we have identified with scRNA-seq.

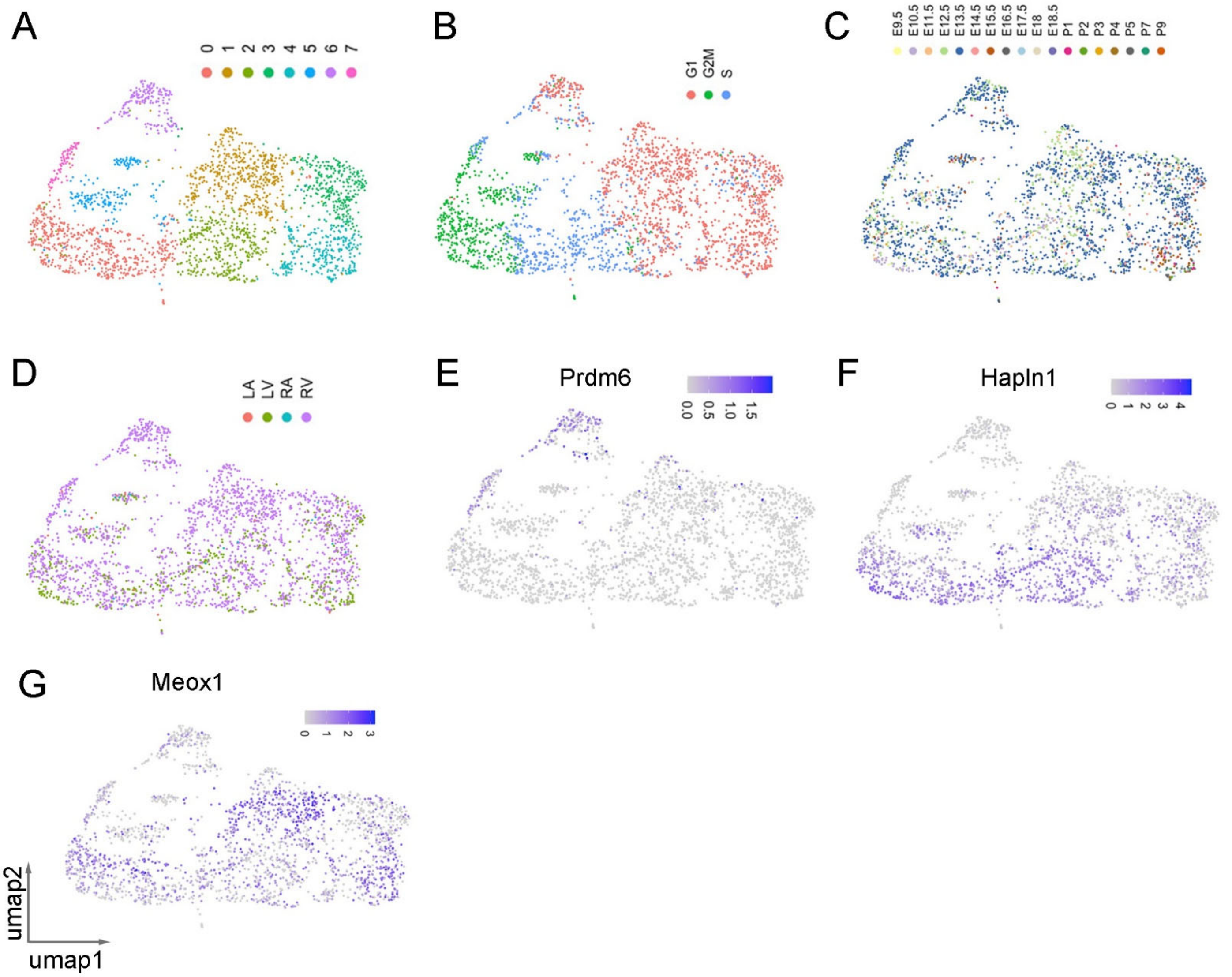


Supplementary Fig S2: Integrative analysis of the valve interstitial cells from CD1 and C57BL/6 scRNA-seq datasets. (A-D) UMAP plots of the VICs labeled by cluster, cell cycle phases, stage, and zone. (E-G) UMAP plots showing the expression of representative VIC genes.

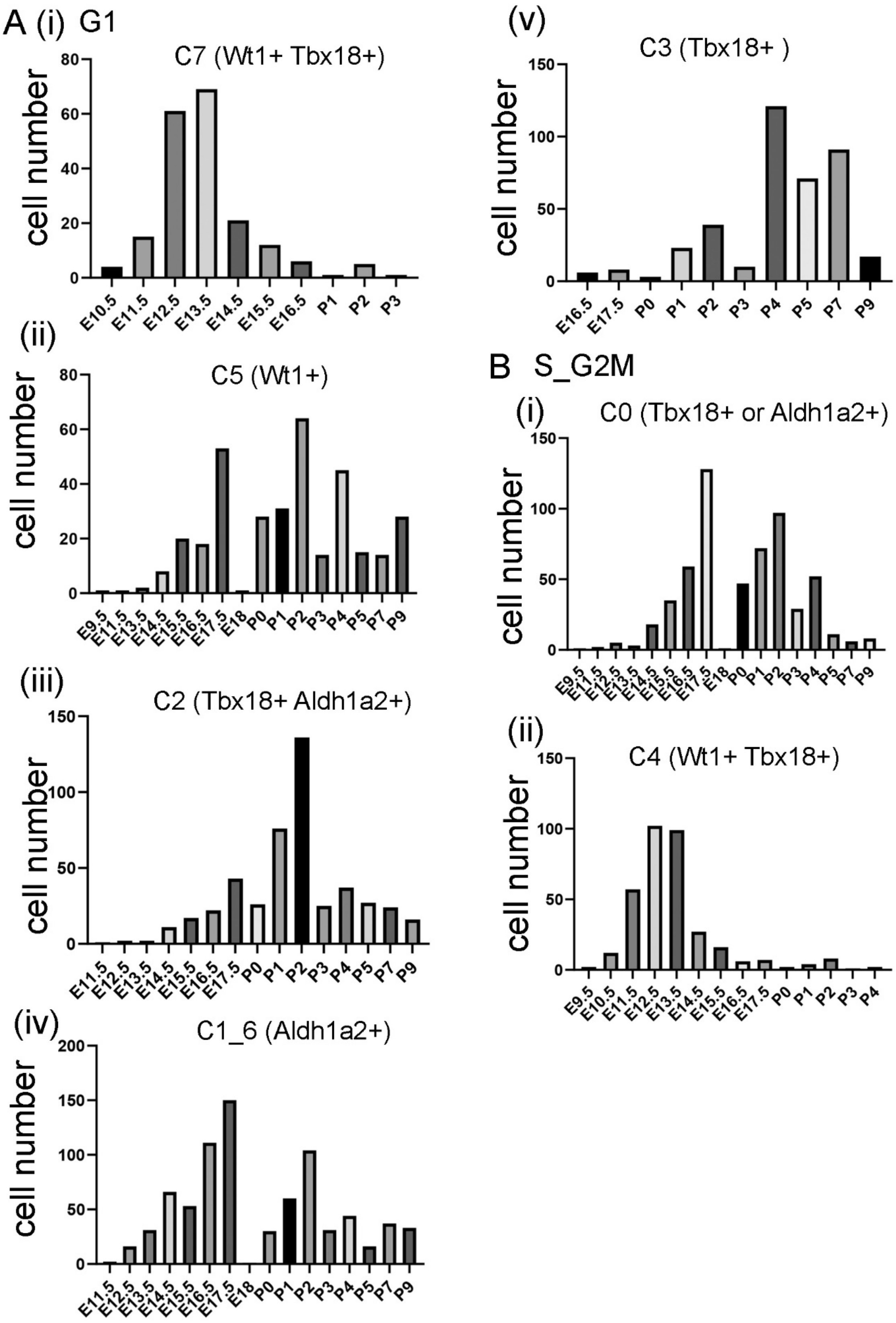


Supplementary Fig S3: Stage analysis of the cells in each main FB subpopulation. (Ai-v) The stage distribution of each subpopulation in G1 phased FBs (CD1 mice). (Bi, ii) The stage distribution of each subpopulation in S_G2M phased FBs (CD1 mice).

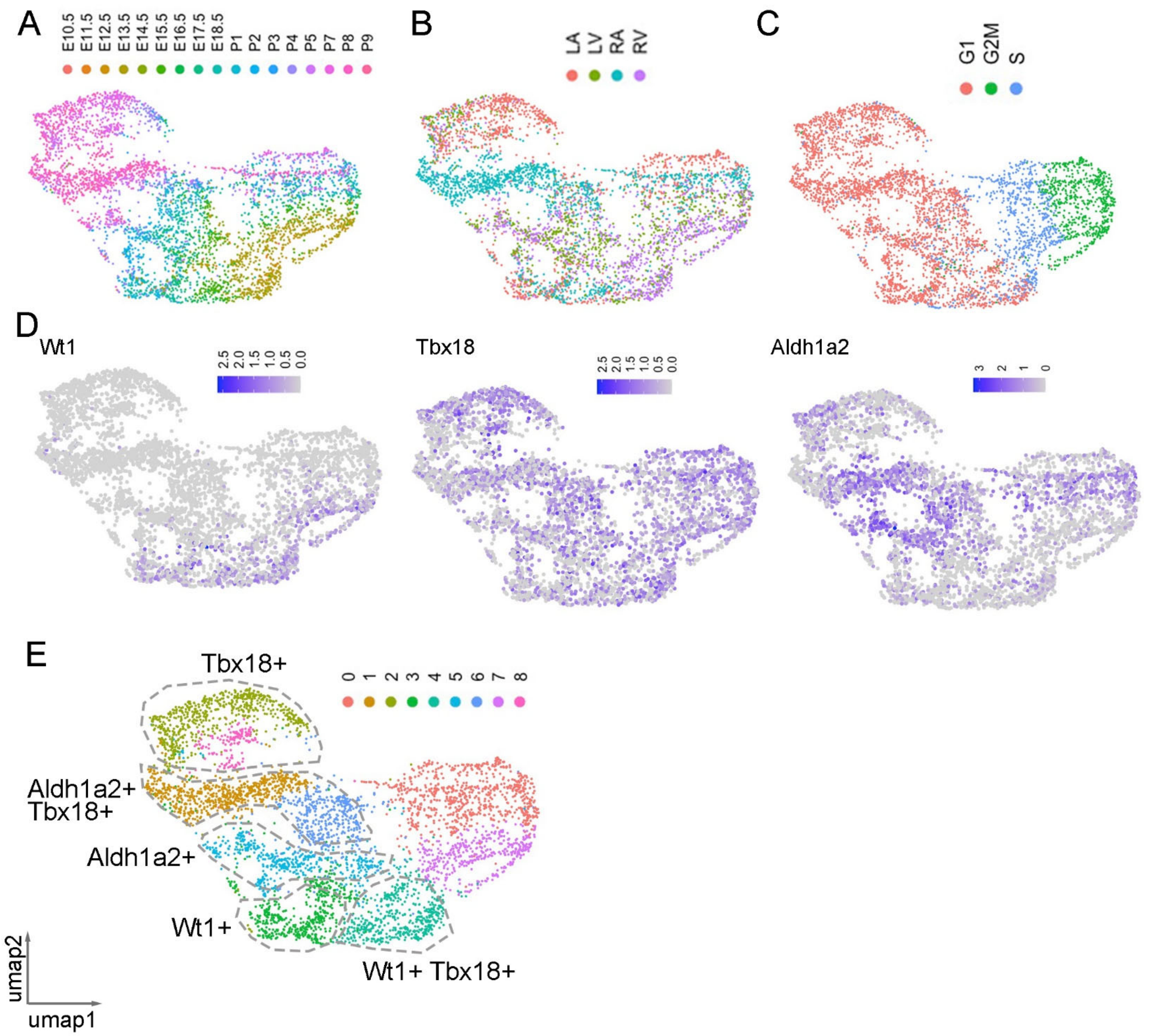


Supplementary Fig S4: The subpopulations in C57BL/6 main FBs. (A-C) UMAP plots of main population of FBs labeled by stage, zone, and cell cycle phases. (D) UMAP plots showing the expression of representative epicardial lineage genes *Wt1*, *Tbx18*, and *Aldh1a2*. (E) The subpopulations within C57BL/6 main FBs.

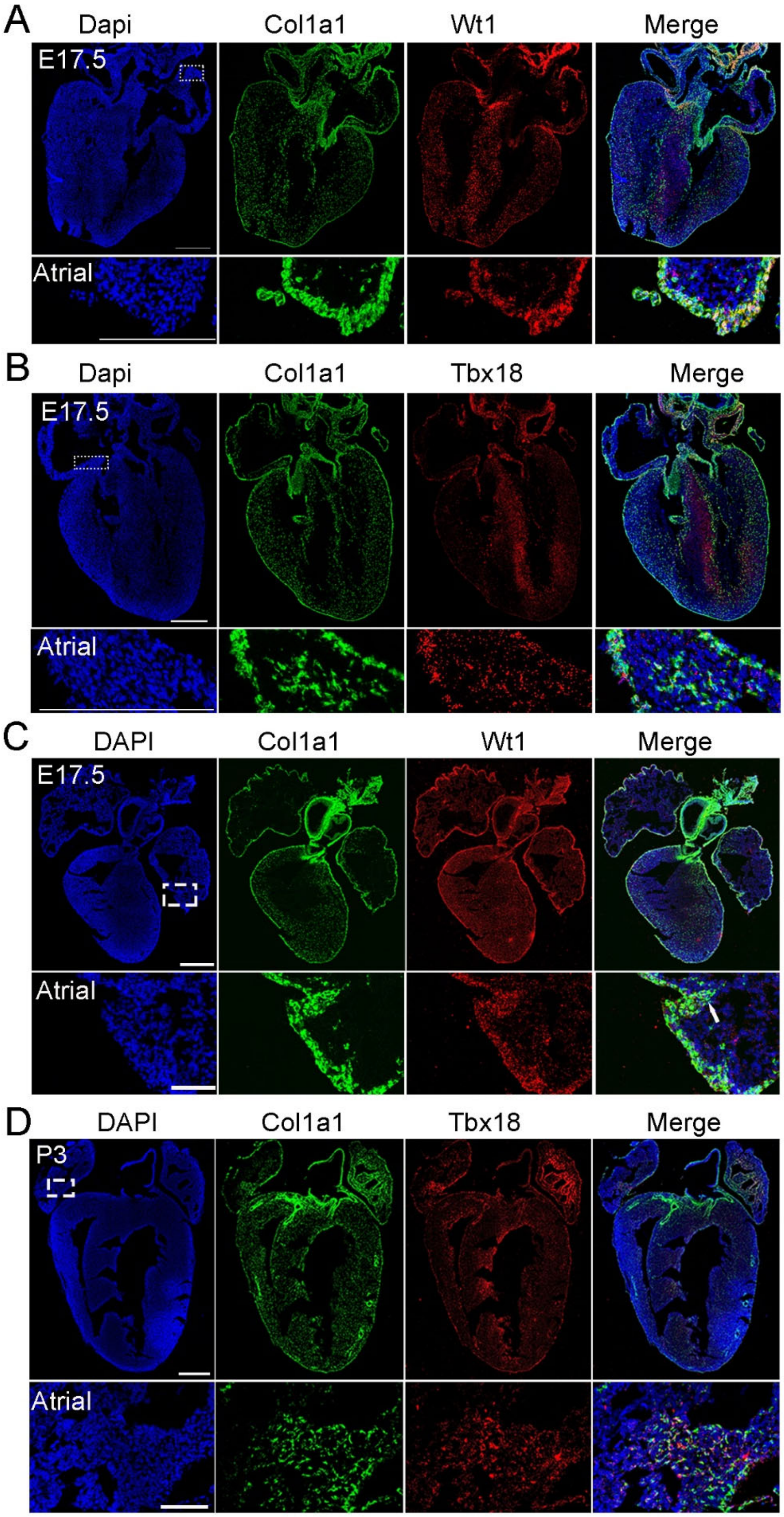


Supplementary Fig S5: Confirmation of FB subpopulations through RNA staining. (A) Co-staining of *Col1a1* and *Wt1* with RNAScope at E17.5. (B) Co-staining of *Col1a1* and *Tbx18* with RNAScope at E17.5. (C) Co-staining of *Col1a1* and *Wt1* with PLISH at E17.5. (D) Co-staining of *Col1a1* and *Tbx18* with PLISH at P3. Scale bar=500μm in the whole heart sections; Scale bar=250μm in the zoomed sections in A and B; Scale bar=100μm in the zoomed section in C and D.

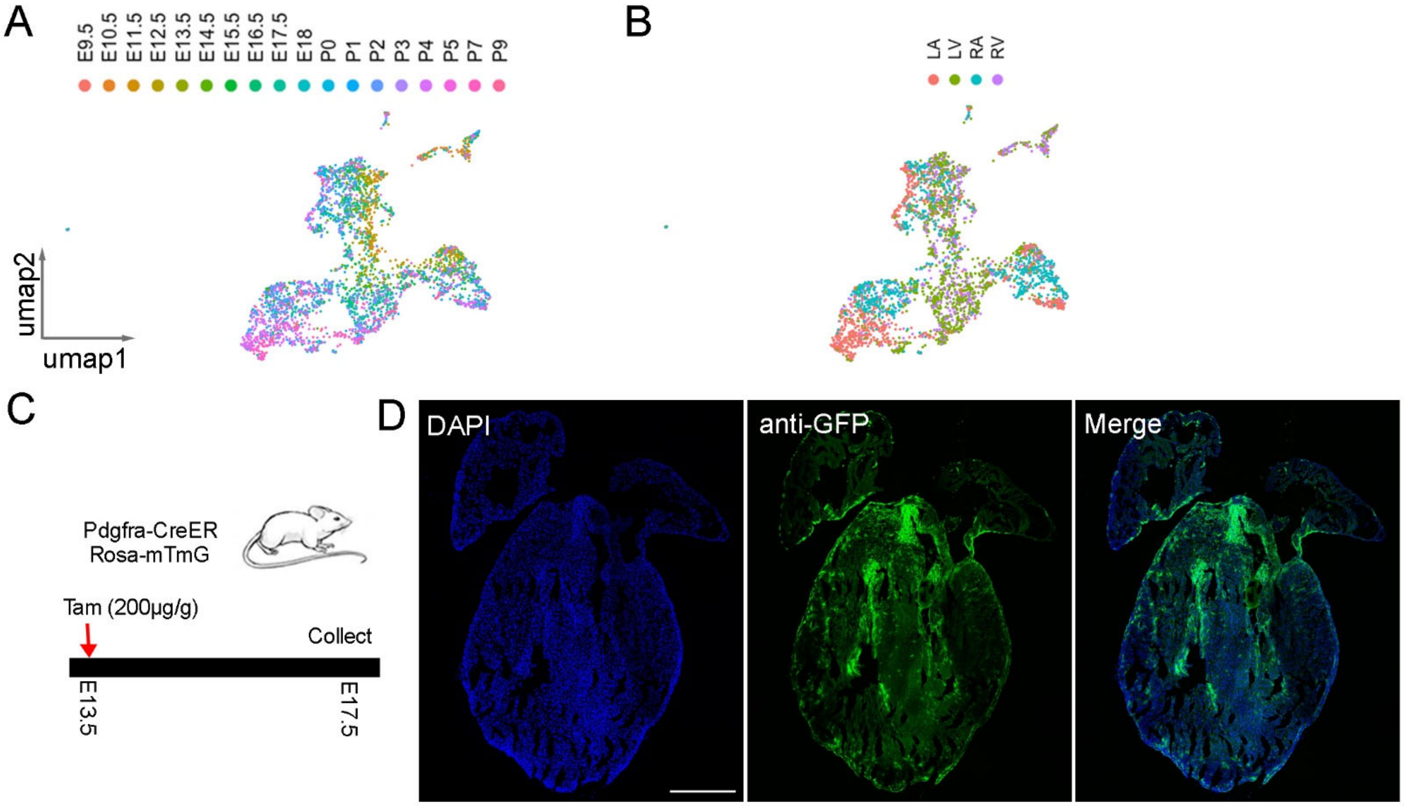


Supplementary Fig S6: (A, B) UMAP plots of FBs in CD1 mice labeled by stage and zone. (C, D) Lineage analysis of Pdgfra-CreER+/-; mTmG+/- mouse hearts with 200ug/g of tamoxifen treatment at E13.5. Scale bar=500μm.

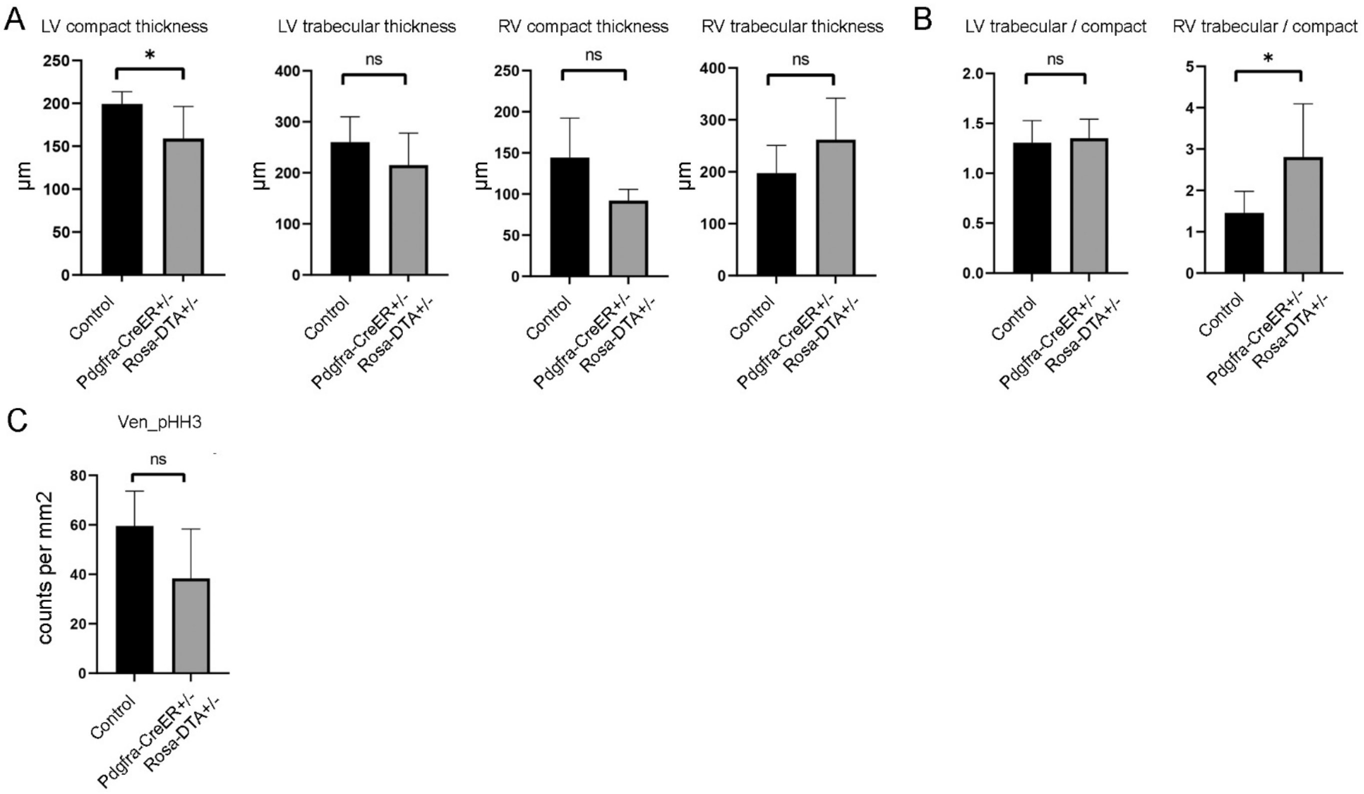


Supplementary Fig S7: Quantification of the defects in control and ablated hearts with one dose of tamoxifen treatment at E15.5. (A) Quantification of the compact and trabecular myocardium thickness in LV and RV. (B) Quantification of the ratio of trabecular to compact myocardium in LV and RV. (C) Quantification of pHH3 positive cells in control and ablated ventricles. * represents $p<0.05$; ** represents $p<0.01$.

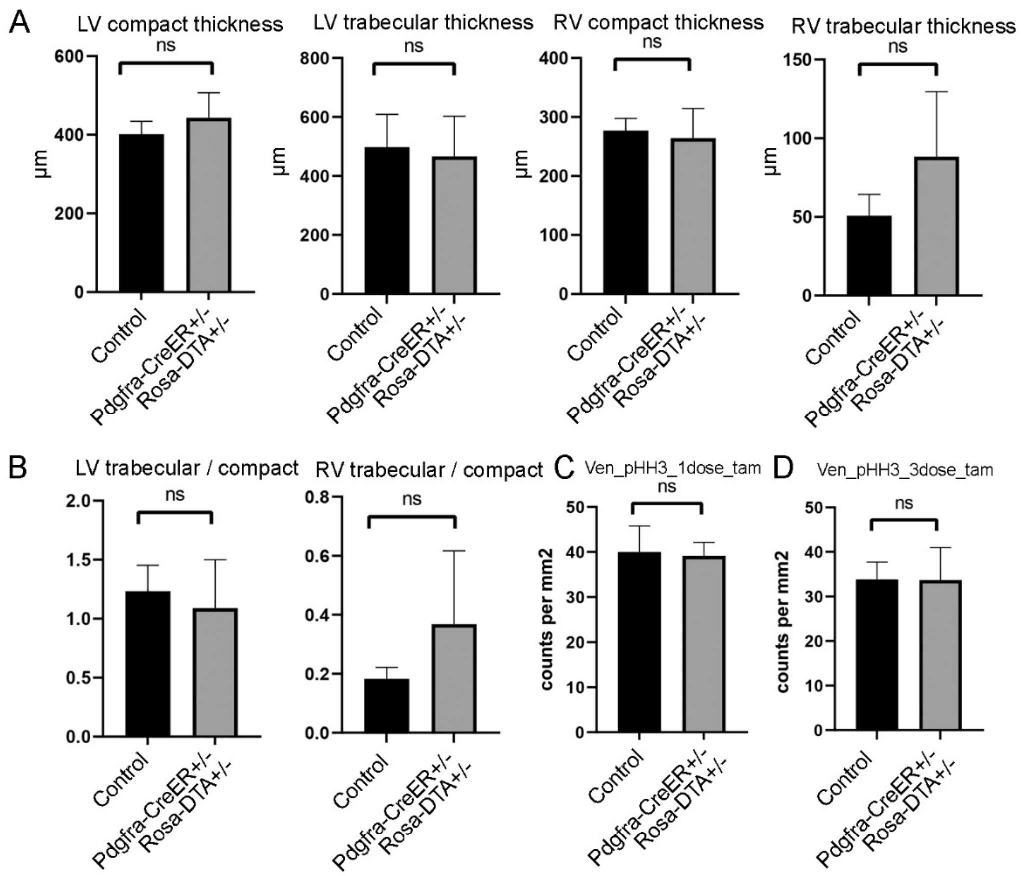


Supplementary Fig S8: Quantification of the defects in control and ablated hearts at neonatal stage. (A) Quantification of the compact and trabecular myocardium thickness in LV and RV after one dose of tamoxifen treatment. (B) Quantification of the ratio of trabecular to compact myocardium in LV and RV after one dose of tamoxifen treatment. (C) Quantification of pHH3 positive cells in ventricular in control and ablated hearts after one dose of tamoxifen treatment. (D) Quantification of pHH3 positive cells in ventricular in control and ablated hearts after three doses of tamoxifen treatments. * represents $p<0.05$; ** represents $p<0.01$.

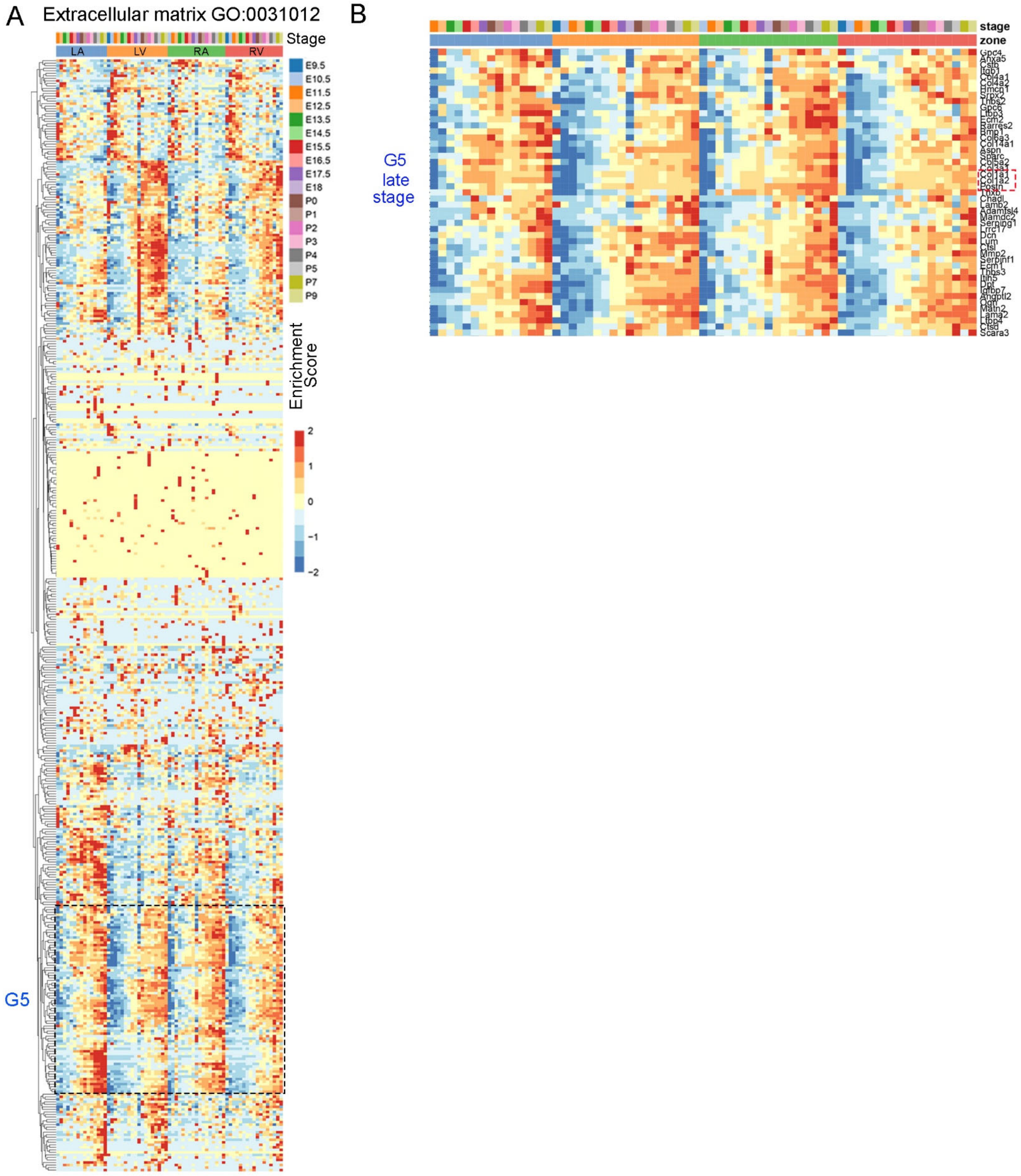


Supplementary Fig S9: (A, B) The expression pattern of a group of genes (G5) that displayed expression in all four chambers of FBs (CD1 mice) at late embryonic and neonatal stages.

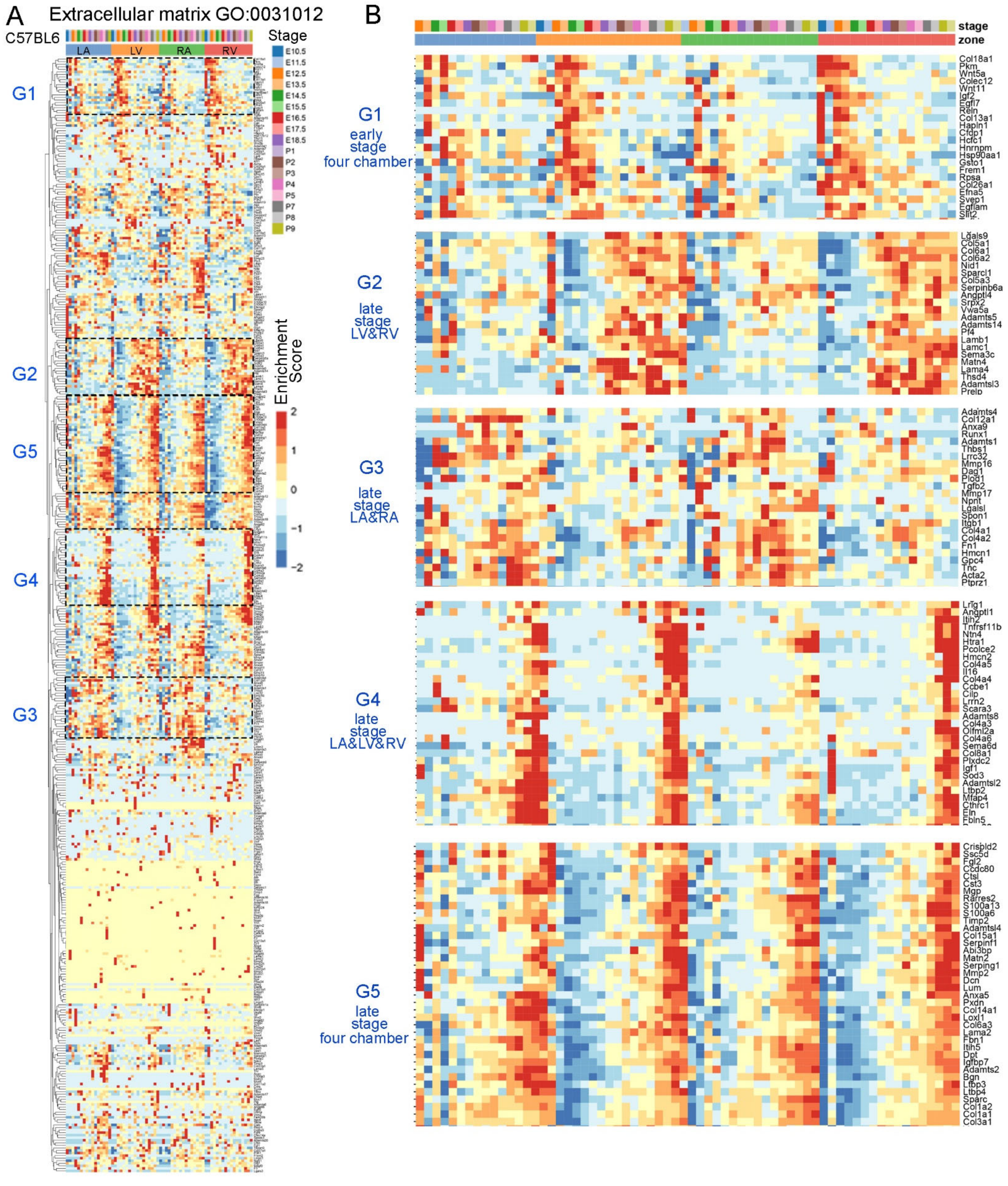


Supplementary Fig S10: The expression pattern analysis of extracellular matrix genes in C57BL/6 FBs. (A) Unsupervised clustering analysis of extracellular matrix genes expression in C57BL/6 FBs. (B) The groups of genes that display stage or zone-specific expression pattern.

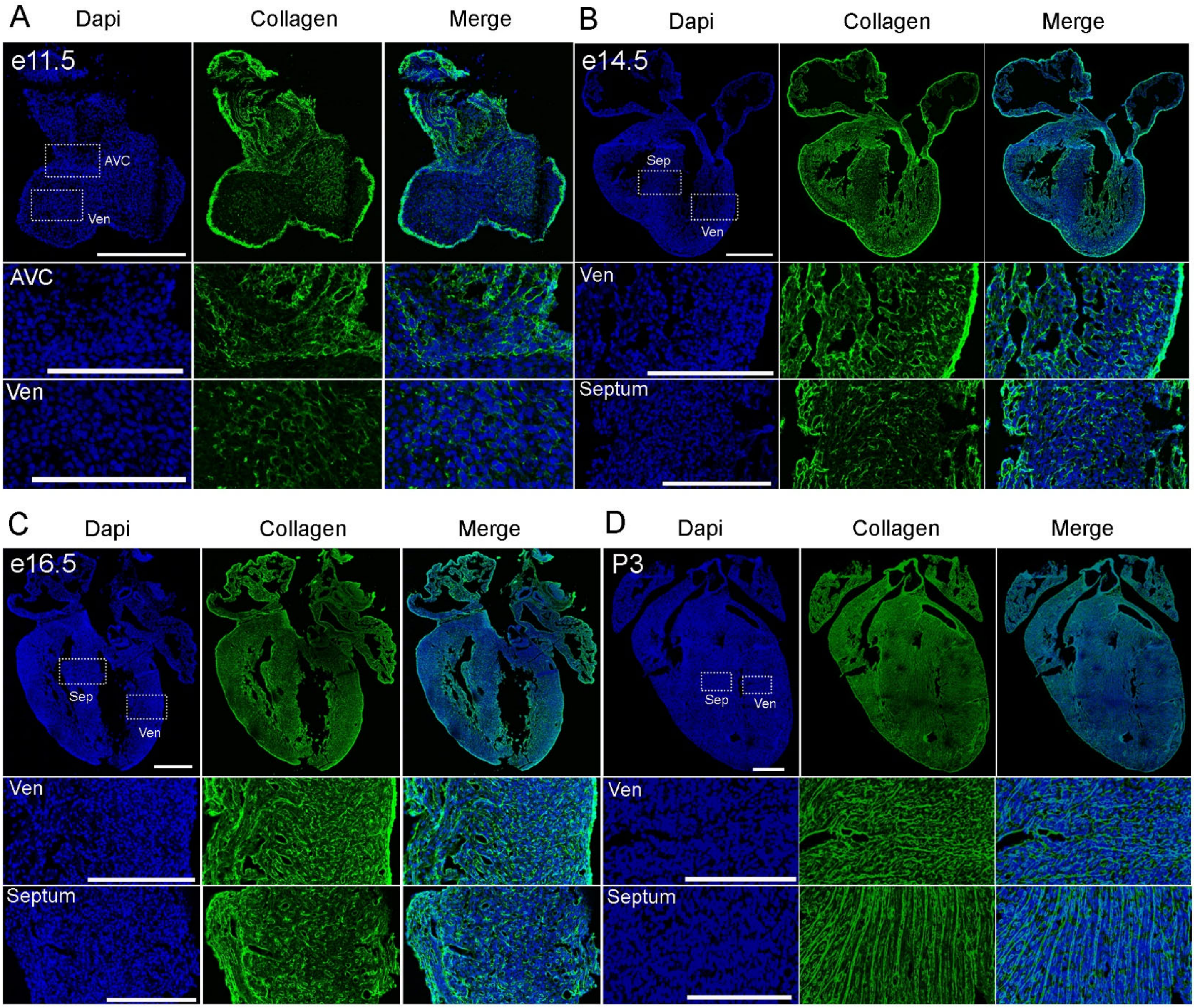


Supplementary Fig S11: Staining analysis of collagen accumulation in different stages of hearts using collagen hybridizing peptide. (A-D) Collagen expression pattern in CD1 mouse hearts at E11.5, E14.5, E16.5, and P3. Scale bar=500um and 150um in the whole heart sections and enlarged sections, respectively.

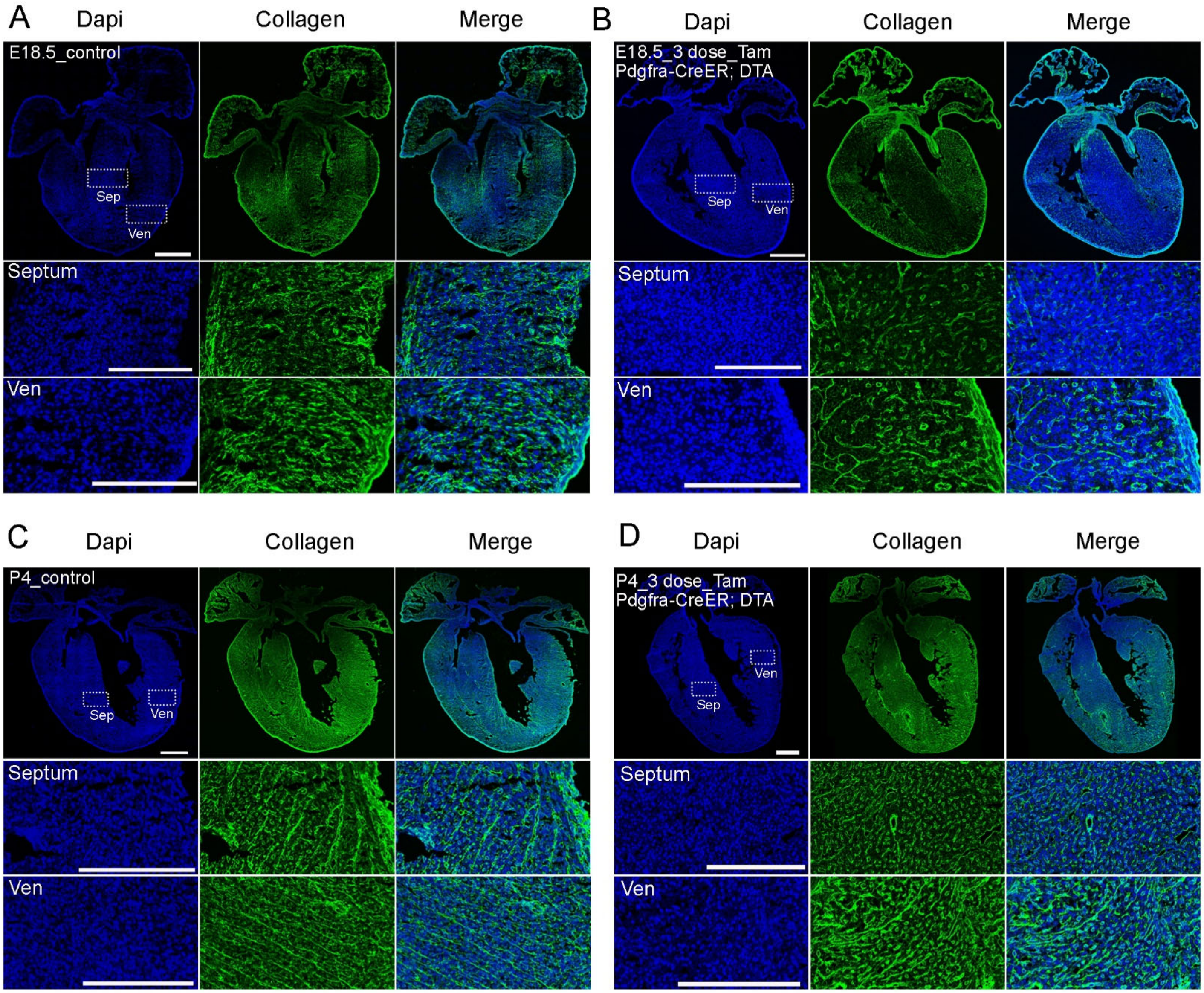


Supplementary Fig S12: In situ analysis of collagen accumulation in control and ablated hearts. (A) Collagen accumulation in control hearts at E18.5. (B) Collagen accumulation in Pdgfra-CreER;Rosa-DTA ablated hearts. Three doses of tamoxifen were given at E15.5, E16.5, and E17.5. (C) Collagen accumulation in control hearts at P4. (D) Collagen accumulation in Pdgfra-CreER;Rosa-DTA ablated hearts. Three doses of tamoxifen were given at P1, P2, and P3. Scale bar=500μm and 150μm in the whole heart sections and enlarged sections, respectively.

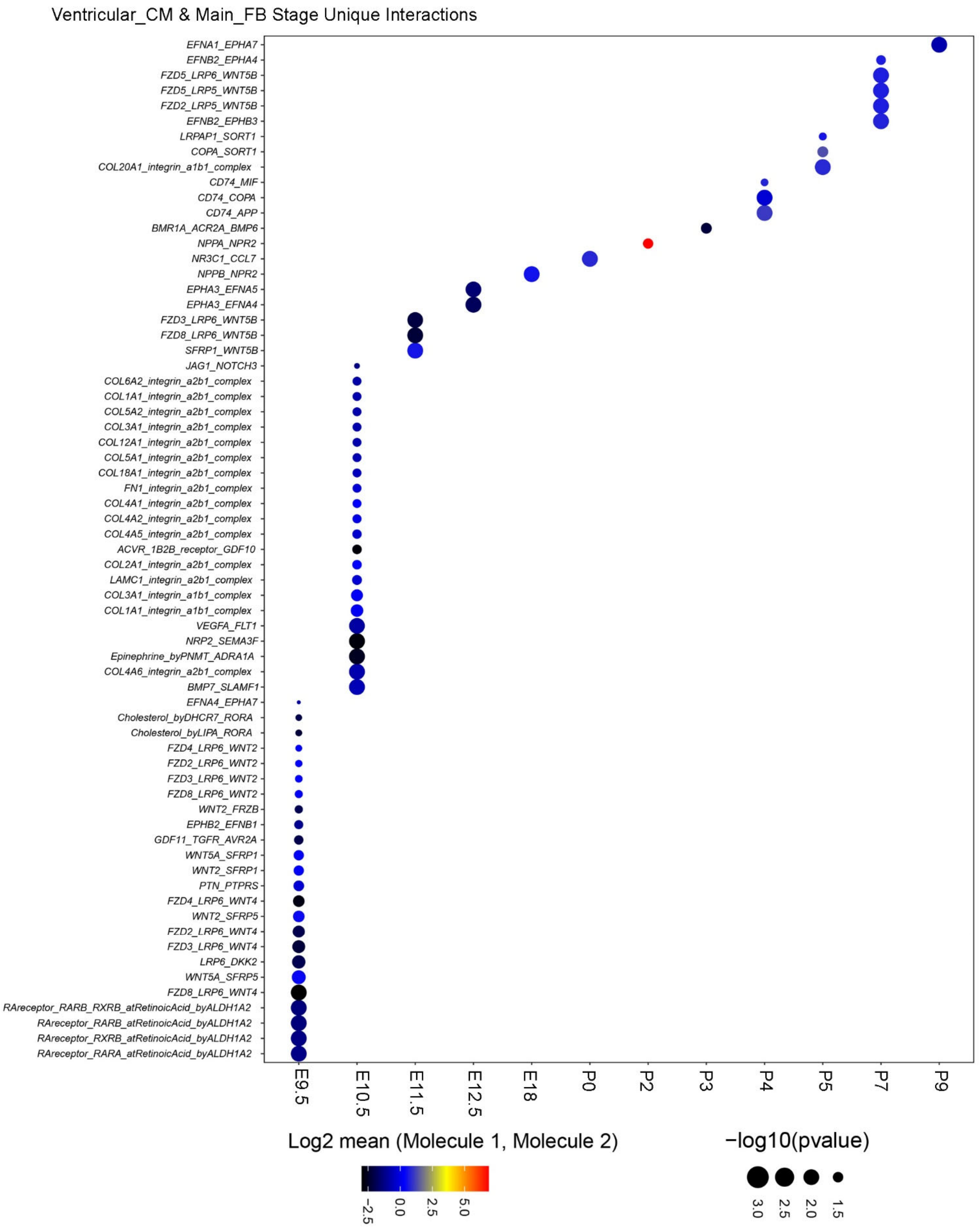


Supplementary Fig S13: The stage-unique ligand receptor interactions between the main population of fibroblast and ventricular CMs.

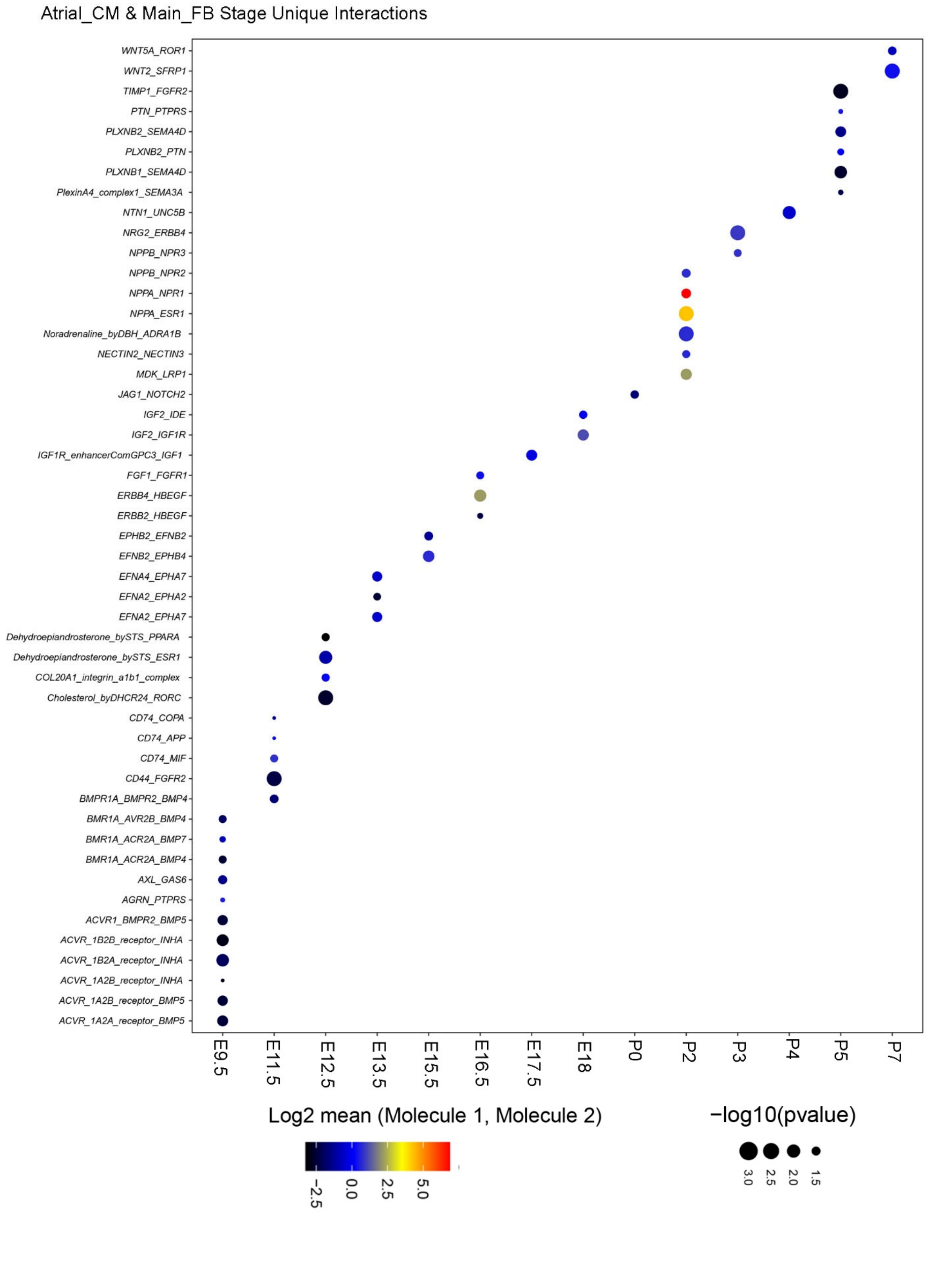


Supplementary Fig S14: The stage-unique ligand receptor interactions between the main population of fibroblast and atrial CMs.
