## Supplemental Tables for "Heterogeneity and Functional Analysis of Cardiac Fibroblasts in Heart Development": TableS1_PLISH probes.docx

| **Name** | **Sequence** |
| --- | --- |
| cy5-mDcn-Right-1 | atgtattttcacgaccttttTTATACGTCGAGTTGAACGTCGTAACA |
| cy5-mDcn-left-1 | TAGCGCTAACAACTTACGTCGTTATGAatccgggtatttgccacag |
| cy5-mDcn-Right-2 | CcaaacccagatcagaacacTTATACGTCGAGTTGAACGTCGTAACA |
| cy5-mDcn-left-2 | TAGCGCTAACAACTTACGTCGTTATGTgcaccactcgaagatgaca |
| cy5-mDcn-Right-3 | CtcttcagtccctggaaggcTTATACGTCGAGTTGAACGTCGTAACA |
| cy5-mDcn-left-3 | TAGCGCTAACAACTTACGTCGTTATGtccgttttcaatcccagagt |
| cy5-mDcn-Right-4 | atgaggaacattggccagacTTATACGTCGAGTTGAACGTCGTAACA |
| cy5-mDcn-left-4 | TAGCGCTAACAACTTACGTCGTTATGtgccattctccataacggtg |
| cy5-mDcn-Right-5 | attttgttgttttgcaggtcTTATACGTCGAGTTGAACGTCGTAACA |
| cy5-mDcn-left-5 | TAGCGCTAACAACTTACGTCGTTATGtagcaaggttgtgtcgggtg |
| Cy5-mAldh1a2-Right-6 | ccgccatttagggattccatTTATACGTCGAGTTGAACGTCGTAACA |
| Cy5-mAldh1a2-left-6 | TAGCGCTAACAACTTACGTCGTTATGagttgcaagagttgccctgt |
| Cy5-mAldh1a2-Right-7 | ggtttgatgaccacggtgttTTATACGTCGAGTTGAACGTCGTAACA |
| Cy5-mAldh1a2-left-7 | TAGCGCTAACAACTTACGTCGTTATGaccacagcacaatgcgggag |
| Cy5-mAldh1a2-Right-8 | ggcccataccctggcaggatTTATACGTCGAGTTGAACGTCGTAACA |
| Cy5-mAldh1a2-left-8 | TAGCGCTAACAACTTACGTCGTTATGattgacgactccgggtggaa |
| Cy5-mAldh1a2-Right-9 | caaaggggctctgcgcatttTTATACGTCGAGTTGAACGTCGTAACA |
| Cy5-mAldh1a2-left-9 | TAGCGCTAACAACTTACGTCGTTATGaaggcattgtaacaattgat |
| Cy5-mAldh1a2-Right-10 | cacggtctttacttctgaatTTATACGTCGAGTTGAACGTCGTAACA |
| Cy5-mAldh1a2-left-10 | TAGCGCTAACAACTTACGTCGTTATGactcccgtaagccaaactca |
| Cy5-mAldh1a2-Right-11 | aaacggtattcacccaggttTTATACGTCGAGTTGAACGTCGTAACA |
| Cy5-mAldh1a2-left-11 | TAGCGCTAACAACTTACGTCGTTATGagagactggcttcgatttgg |
| Cy5-mAldh1a2-Right-12 | ttgaagggagctagctggttTTATACGTCGAGTTGAACGTCGTAACA |
| Cy5-mAldh1a2-left-12 | TAGCGCTAACAACTTACGTCGTTATGaatcgtgtgttcacatgggt |
| cy5-mCol1a1-Right-1 | CCACCCCTTCACAGAGATGTTTATACGTCGAGTTGAACGTCGTAACA |
| cy5-mCol1a1-left-1 | TAGCGCTAACAACTTACGTCGTTATGAGCACCTTTGATACCAAACT |
| cy5-mCol1a1-Right-2 | ACACAATTGCACTGAGGAATTTATACGTCGAGTTGAACGTCGTAACA |
| cy5-mCol1a1-left-2 | TAGCGCTAACAACTTACGTCGTTATGAGAACGGTCTCTCCCACCCA |
| cy5-mCol1a1-Right-3 | CATGGAGATGCCAGATGGTTTTATACGTCGAGTTGAACGTCGTAACA |
| cy5-mCol1a1-left-3 | TAGCGCTAACAACTTACGTCGTTATGAGGTTCCTTCAACAGTCCAA |
| cy5-mCol1a1-Right-4 | GACTTATACCCACATAGGTCTTATACGTCGAGTTGAACGTCGTAACA |
| cy5-mCol1a1-left-4 | TAGCGCTAACAACTTACGTCGTTATGTTCAAGCAAGAGGACCAAGC |
| cy5-mCol1a1-Right-5 | GCCCCAAGTTCCGGTGTGACTTATACGTCGAGTTGAACGTCGTAACA |
| cy5-mCol1a1-left-5 | TAGCGCTAACAACTTACGTCGTTATGTCGTGCAGCCGTCCACAAGG |
| Cy3-mHapln1-Right-1 | GGCAGTGTCACGTTGCCACCTTATACGTCGAGTTGACCGACGTATTG |
| Cy3-mHapln1-left-1 | TATTCGTTCGAACTTACGTCGTTATGTCGGTGAGAGAAGACCTTGG |
| Cy3-mHapln1-Right-2 | CATCACCTTGATACTCTGGTTTATACGTCGAGTTGACCGACGTATTG |
| Cy3-mHapln1-left-2 | TATTCGTTCGAACTTACGTCGTTATGAATCTTGAAGTCTCGAAAGG |
| Cy3-mHapln1-Right-3 | CTGCCAGCAACCAAATTGGTTTATACGTCGAGTTGACCGACGTATTG |
| Cy3-mHapln1-left-3 | TATTCGTTCGAACTTACGTCGTTATGATCTTATTTGAGTCTACTAA |
| Cy3-mHapln1-Right-4 | CATGACACCAGCATGCTTTTTTATACGTCGAGTTGACCGACGTATTG |
| Cy3-mHapln1-left-4 | TATTCGTTCGAACTTACGTCGTTATGAGGAGTACTGAAATCAAGAA |
| Cy3-mHapln1-Right-5 | TTTAAATATTGACCTTCTTCTTATACGTCGAGTTGACCGACGTATTG |
| Cy3-mHapln1-left-5 | TATTCGTTCGAACTTACGTCGTTATGTAATTGGGTGGCTTACTGAT |
